# Assistive sensory-motor perturbations influence learned neural representations

**DOI:** 10.1101/2024.03.20.585972

**Authors:** Pavithra Rajeswaran, Alexandre Payeur, Guillaume Lajoie, Amy L. Orsborn

## Abstract

Task errors are used to learn and refine motor skills. We investigated how task assistance influences learned neural representations using Brain-Computer Interfaces (BCIs), which map neural activity into movement via a decoder. We analyzed motor cortex activity as monkeys practiced BCI with a decoder that adapted to improve or maintain performance over days. Over time, task-relevant information became concentrated in fewer neurons, unlike with fixed decoders. At the population level, task information also became largely confined to a few neural modes that accounted for an unexpectedly small fraction of the population variance. A neural network model suggests the adaptive decoders directly contribute to forming these more compact neural representations. Our findings show that assistive decoders manipulate error information used for long-term learning computations like credit assignment, which informs our understanding of motor learning and has implications for designing real-world BCIs.

---

We learn new motor skills by incrementally minimizing movement errors [1]. This process involves both widely distributed computations [2] and targeted neurophysiological changes [3, 4, 5]. Such coordinated circuit changes might be mediated by a poorly understood “credit assignment” process where movement errors lead to adjustment in selected neural structures [6].

Intracortical brain-computer interfaces (BCIs) provide a powerful framework to study how motor errors shape neural changes [7, 8]. BCIs define a mapping (“decoder”; Fig. 1A) between neural activity and movement [9, 10, 11] to causally interrogate learning. Perturbations of well-learned BCI mappings, analogous to perturbations of natural movement [12, 13], have been used to probe the neural mechanisms of short-term adaptation by introducing task errors [14, 15]. Alternately, novel BCI mappings learned over multiple practice sessions can be used to study neural mechanisms related to skill formation [16, 17, 18, 19]. BCI skill learning studies often use decoders that provide limited performance, requiring the brain to reduce task errors through changes in neural activity. In contrast, application-driven BCIs often manipulate decoders to *improve* or *maintain* performance over time (Fig. 1A,B) [11, 20, 21, 22, 23, 24, 25, 26, 27, 28]. These assistive interventions perturb the mapping between neural activity and movement without introducing task errors and potentially reducing them, thus providing an opportunity to study how errors influence skill learning mechanisms.

**Figure 1.**
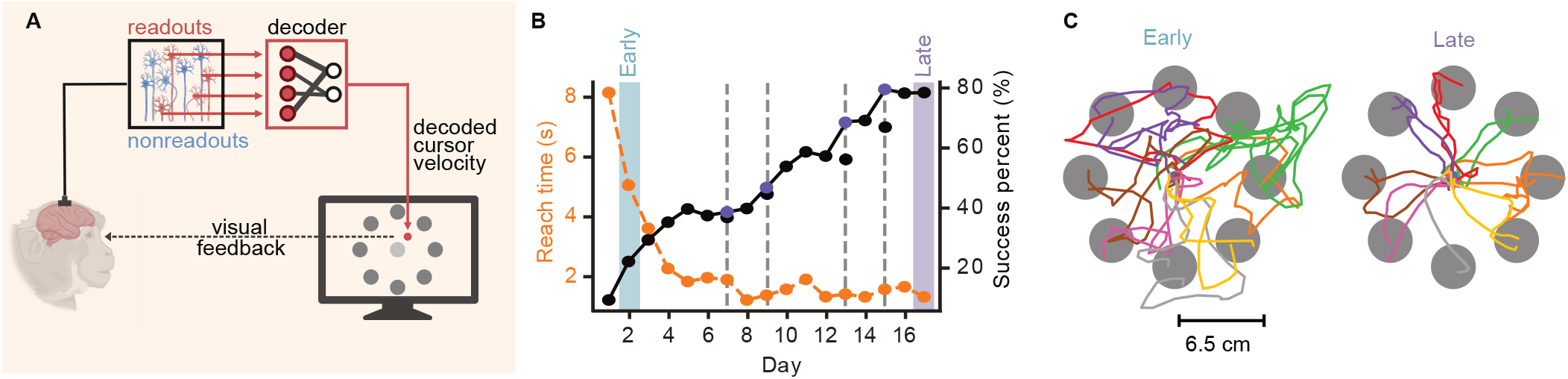
Experimental setup and example behavior. **(A)** Closed-loop BCI setup. Multi-unit activities from readouts (red) and nonreadouts (blue) were recorded in a monkey’s motor cortex. Only the readout activity was decoded to control a cursor in a 2D center-out reaching task with visual feedback. (Monkey and neuron icons from BioRender.) **(B)** Mean reach time (dashed orange line) and success percentage (solid black line) across days for an example learning series (monkey J). Early (blue) and late (purple) training phases are indicated. Vertical grey dashed lines indicate decoder adaptation, with purple dots showing performance gains from decoder adaptation. **(C)** Early (left) and late (right) cursor trajectories for the series in panel B.

BCI skill learning alters both single-neuron and population activity. Practice with the same decoder over several days refines performance, stabilizes neural representations [16] and improves coordination among neurons [29, 30], which is consistent with reduced neuronal variability in natural motor learning [31]. Some changes are targeted to specific neurons: neurons used as direct “readouts” of behavior are modulated differently than “nonreadout” neurons, which may participate in computations but do not drive movement directly (Fig. 1B). Preferential modulation of readouts with extended practice is consistent with the possibility that credit assignment computations contribute to skill learning [30, 32, 33, 34, 35, 36]. Population-level changes also appear to be primarily subserved by readouts [30].

Long-term learning may also occur during extended practice with assistive BCI decoders. Assistive decoders adapt the neural-movement mapping to reduce errors, and also help compensate for unstable measurements by modifying which neural units are part of the readout ensemble [28] (Fig. 1B). Despite this dynamic mapping, adaptive decoders can help produce skillful control of BCI over time[11, 21, 28, 36], and reduce the amount of neural changes needed, such as shifts in neuronal tuning [28]. Yet, longitudinal adaptive BCI experiments consistently report performance improvements beyond what would be expected from decoder adaptation alone [11, 21, 28, 36, 37, 38, 39], consistent with long-term skill learning processes taking place.

It is unclear how decoder interventions that manipulate errors impact learning and the resulting neural representations. We hypothesized that adaptive BCIs might alter feedback that contributes to the targeted neural changes that occur during skill learning. Better understanding this process could inform BCI applications that widely use these manipulations and provide insights into the computations guiding long-term sensorimotor learning. However, examining brain-algorithm interactions is challenging because it requires comparing learning outcomes between different paradigms (fixed vs. adaptive decoders) where variations in task performance, individuals, and training history complicate interpretations. Existing work has therefore focused on performance-based comparisons of different BCI algorithms without examining their impact on neural representations.

We used experimental data from longitudinal BCI learning studies [16, 28] and recurrent neural network (RNN) models to explore how adaptive decoders influence learned neural representations. The experimental data provided insights into real-world neural adaptation, while the RNN model allowed for direct comparisons that are not possible experimentally. Our findings show that learning-mediated changes were targeted to the readouts, consistent with credit assignment computations. But surprisingly, these changes were mostly concentrated in a subset of readouts when decoders adapted. In contrast, fixed-decoder experiments did not show such *compaction* with learning. Our model helped ascertain the direct involvement of decoder adaptation in the compaction process, and offered insight into how adaptive decoders could influence plasticity. Modeling also suggested opportunities to tune brain-decoder interactions to balance performance goals. Finally, data analysis of population activity revealed that learning mainly targeted low-variance components, which are often overlooked by standard methods [40]. Together, our findings provide evidence that adaptive decoders influence neural encoding on multiple levels, shedding new light on how error manipulations shape neural activity during learning.

## Results

Two rhesus macaques (monkeys J and S) learned to perform point-to-point movements with a BCI controlled cursor (Fig. 1A), as previously described [28]. Multi-unit action potentials from microelectrode arrays implanted in the arm area of the primary- and premotor cortices were used as inputs to a Kalman filter decoder that controlled cursor velocity and position. A subset of all recorded units was selected as input to the decoder based on recording properties (e.g., stability over time) and defined the *readout* population (Fig. 1A) (see Methods).

Each monkey learned to control the cursor over several consecutive days, defined as a learning series. We focused on learning series lasting at least four days (*n* = 7 series for monkey J; *n* = 3 for monkey S). Learning series varied in length, initial decoder training methods, and number of decoder updates. Decoder weights were updated via Closed-Loop Decoder Adaptation (CLDA) by realigning cursor velocities produced by readouts toward the target (see Methods and Fig. S1A). CLDA was done at the beginning of a learning series, improving initial performance to provide sufficient cursor control throughout the workspace. CLDA was also done intermittently within series to maintain performance when neural measurements varied. Longer series generally had more decoder adaptation events (Fig. S1D). As previously shown [28], task success rates and reach times improved over days (Fig. 1B, a selected series for monkey J used below in all single-series examples, for consistency; see additional example series in Fig. S1B (monkey S) and Fig. S1E (monkey J)).

We quantified learning within each series by defining an ‘early’ and ‘late’ training day, which corresponded to the first and last day of a learning series with at least 25 successful reaches per target direction, respectively (day 2 and 17 in Fig. 1B). We were primarily interested in characterizing neural representations related to movement direction, and therefore included any trial wherein the target was reached regardless of whether the hold at the target was successful. Improvements in task success rate and reach time were accompanied by more direct and accurate reaches (Fig. 1C, Fig. S1C and [28]). The dataset included series where the amount of CLDA performed on day 1 varied, resulting in varying initial performance, but behavioral improvements consistent with long-term skill learning were observed in all series [28].

### Credit assignment to readout units during long-term learning with adaptive decoders

We first assessed whether adaptive decoders impacted credit assignment in motor cortex by investigating how task-relevant information was distributed between readouts and nonreadouts. Because cursor kinematics changed significantly across days, we instead used an offline classification analysis to quantify how the fixed target identities were encoded. Target identity does not change over days, facilitating cross-day comparisons despite behavioral changes (Fig. 1C). Specifically, each day we fitted a multiclass logistic regression to predict target identity using neural activity aligned on the go-cue (Fig. 2A), evaluating classification accuracies using cross-validation. This classifier accurately predicted target identity per trial, with mean accuracies of 0.78 ± 0.13 (monkey S) and 0.83 ± 0.11 (monkey J) across all days and all learning series.

**Figure 2.**
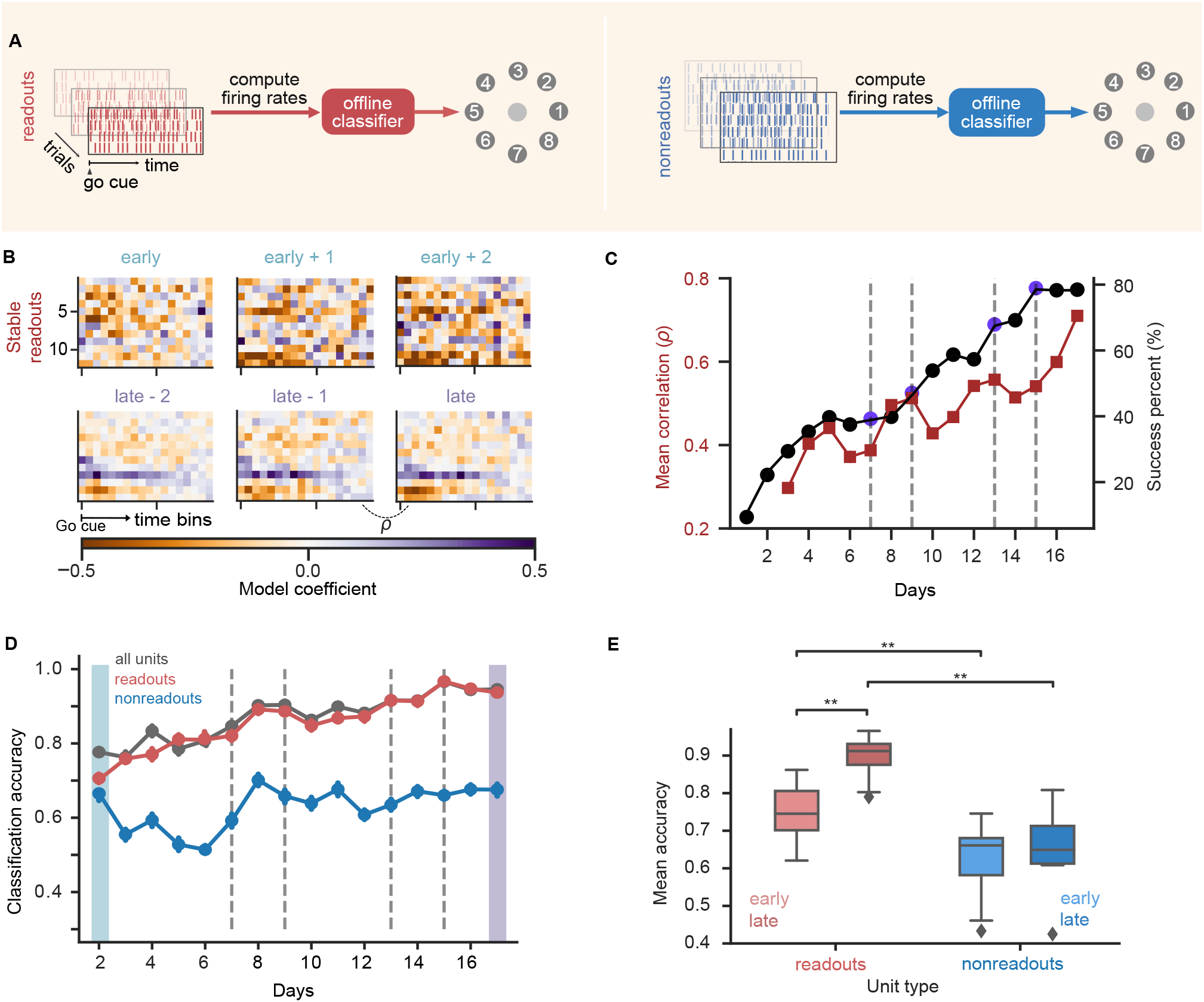
Credit assignment to readout units. **(A)** Offline classification model. **(B-D)** Classification results for the selected series (Fig. 1D). **(B)** Heatmap representation of the classifier weights to predict a single target (target 2) for the stable readout population on multiple days. **(C)** Success percentage (black; reproduced from Fig. 1D) and mean correlation coefficient between classifier weights on consecutive days (dark red) plotted across days. **(D)** Classification accuracy for all units (gray), readout units (red) and nonreadout units (blue). Error bars represent 95% confidence intervals on test accuracy (see Methods). **(E)** Early vs late classification accuracy for readout (shades of red) and nonreadout populations (shades of blue) across all series for both monkeys. (Readout-early vs readout-late : ** *p* = 0.006; readout-early vs nonreadout-early: ** *p* = 0.002; readout-late vs nonreadout-late: ** *p* = 0.002; nonreadout-early vs nonreadout-late: *p* = 0.13 (ns). Two-sided Wilcoxon signed-rank test, *n* = 10.)

The logistic regression model weighted the contribution of each unit’s firing rate across time bins between the go-cue and trial end towards classifying target identity. These weights thus represent an encoding of task information, and are related to parametric analyses of neuron activity such as direction tuning (Fig. S2C). We found that optimized classifier weights varied day to day, but became more similar as learning progressed (Fig. 2B). The correlation coefficient of the weights of stable readouts on consecutive days increased during learning (Fig. 2C), consistent with a progressive stabilization of task encoding representations that mirrors performance improvements (Fig. 1D). These findings are consistent with previous observations that adaptive BCIs lead to stable directional tuning in readout neurons [28].

Target classification accuracy improved during BCI training when using all neurons, paralleling the gains in BCI performance (Fig. 2D). However, this improvement was mostly driven by the readouts (Figs. 2D, E). Classification accuracy was higher for readouts compared to nonreadouts both early and late in training despite large differences in their respective numbers (Supplementary Table 1). Similar results were obtained with other classifiers and with regularization (Fig. S2A). More importantly, only the readouts’ accuracy increased significantly from early to late (Fig. 2E; Fig. S2F for individual animals), showing that learning-related changes in classification accuracy are driven predominantly by changes within the readout population. We observed a similar separation in information encoding between readout and nonreadout populations when analyzing only the time period immediately after the go-cue (0-200 ms), which further controls for potential differences in reach kinematics with learning (Fig. S2G). The same analysis on data during arm movements, in contrast, showed no significant separations between readout and nonreadout populations (Fig. S2D,E), ruling out factors such as variability in neural recording quality and confirming credit assignment was specific to the BCI task [32].

**Table 1.** Parameters of the model. For the arm model, we combined data from Ref. [64] and from the morphology of the subjects in Ref. [28].

| Parameters of the arm model |  |  |  |
| --- | --- | --- | --- |
| *symbol* | *value* | *unit* | *description* |
| $m_1$ | 0.25 | kg | mass of upper arm link |
| $m_2$ | 0.20 | kg | mass of lower arm link |
| $l_1$ | 0.12 | m | length of upper arm link |
| $l_2$ | 0.16 | m | length of lower arm link |
| $d_1$ | 0.075 | m | distance from center upper arm to center of mass |
| $d_2$ | 0.075 | m | distance from center lower arm to center of mass |
| $I_1$ | 0.0018 | kg·m<sup>2</sup> | moment of inertia of upper arm |
| $I_2$ | 0.0012 | kg·m<sup>2</sup> | moment of inertia of lower arm |
| $\tau_f$ | 40 | ms | time constant connecting torques and controls |
| Parameters of the network dynamics |  |  |  |
| $N$ | 100 | — | number of units |
| $\alpha_v$ | 0.2 | — | reciprocal of time constant |
| $\sigma$ | 0.02 | — | noise intensity |
| Parameters of the objective |  |  |  |
| $\delta_p$ | 0.01 | m | scaling factor for position |
| $\delta_v$ | 0.02 | m/s | scaling factor for velocity |
| $\delta_a$ | 0.08 | m/s<sup>2</sup> | scaling factor for acceleration |
| $\gamma_v$ (arm) | 0.25 | — | velocity cost hyperparameter for arm control |
| $\gamma_v$ (BCI) | 0.1 | — | velocity cost hyperparameter for BCI control |
| $\gamma_a$ | 0.05 | — | acceleration cost hyperparameter for arm control |
| $\lambda_u$ | 0.005 | N<sup>-2</sup>·m<sup>-2</sup> | effort penalty hyperparameter |
| Parameters of the learning algorithm |  |  |  |
| $\alpha_R$ | 0.3 | — | relative weight factor for reward traces |
| — | $2 \times 10^{-5}$ | — | learning rate |

Finally, we examined the temporal relationship between decoder interventions and credit assignment. We compared classification accuracies pre- and post-decoder change events and observed no significant changes (Fig. S2H). This suggests that credit assignment computations were not immediately impacted by decoder changes and instead occurred over longer timescales of practice. Together our results show that assistive decoder perturbations lead to the formation of a stable BCI task neural encoding that is primarily supported by readout units, similar to BCIs with fixed decoders [16, 32].

### Compaction of neural representations with long-term learning alongside decoder adaptation

Our classifier analysis revealed a striking pattern: within a given readout ensemble, a small number of readouts carried the majority of weight for target prediction after learning (Fig. 2B, bottom row). This suggests further credit assignment within the readout population that has not been reported in past BCI studies. Does long-term learning lead certain readouts to become more important? We explored this question by conducting a rank-ordered neuron adding curve (NAC) analysis [41]. This technique quantifies how classification accuracy improves as neurons are sequentially added to a decoding model, ranked from most to least predictive (Fig. 3A).

**Figure 3.**
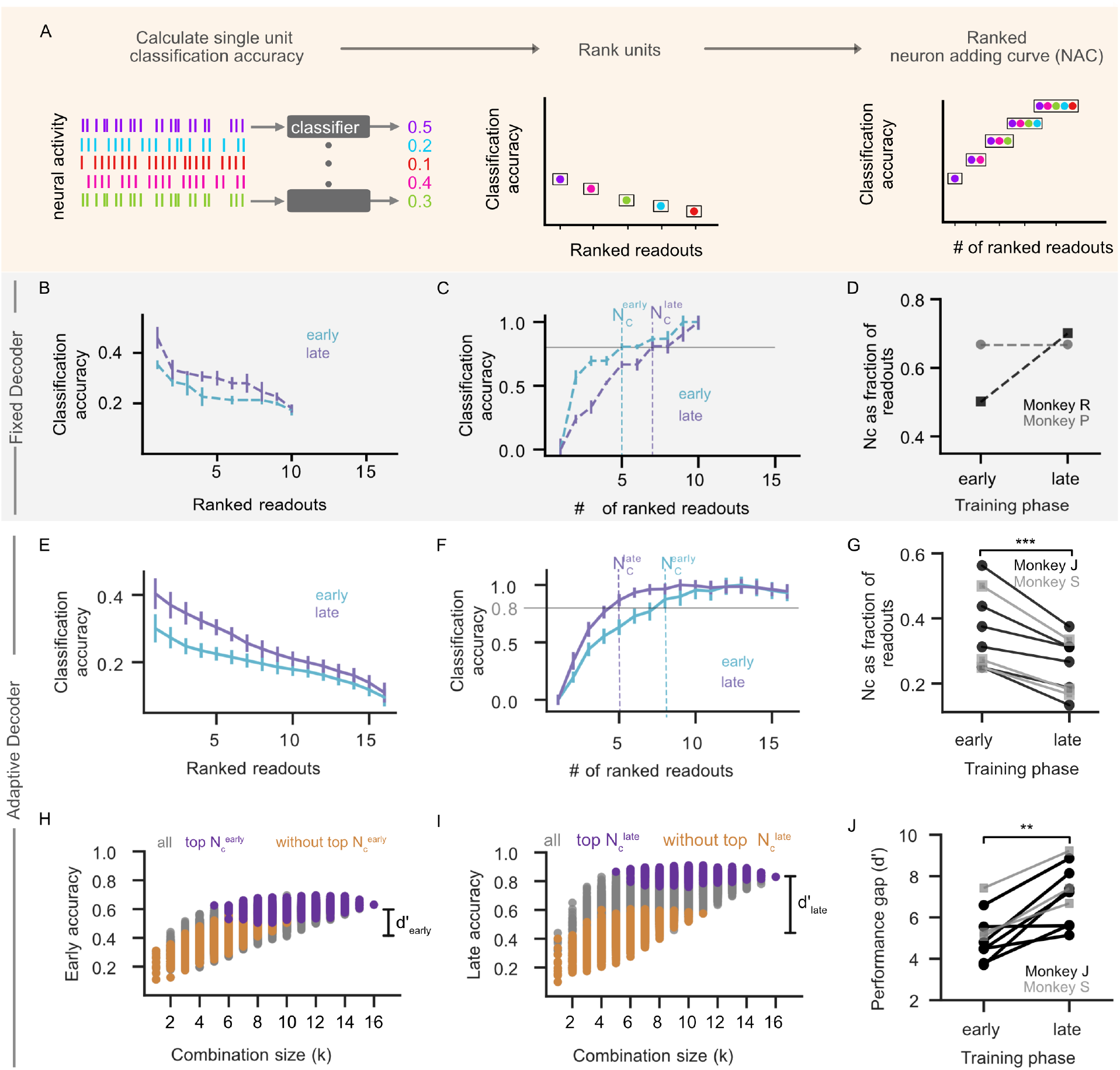
Learning leads to compaction of neural representations: **(A)** Ranked Neuron Adding Curve (NAC) analysis. We first estimated classification accuracies of individual units (left), ranked units based on their classification power (middle) and constructed NACs by incrementally adding the next best unit (right) (see Methods). **(B)** Ranked classification accuracy of individual units on early (cyan) and late (purple) epoch from fixed decoder dataset (example from monkey R). For the fixed decoder, early and late refer to epochs, not days, in contrast to the adaptive decoder. Epochs consist of a constant number of trials, consistent with previous analyses [30]. **(C)** Ranked NAC on early (cyan) vs late (purple) epoch for fixed decoder series from Monkey R. Accuracy is normalized to each epoch’s peak performance. Vertical dashed lines indicate number of units required to reach 80% of peak accuracy (*N*). **(D)** Comparison of 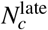 and 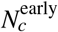 from 2 monkeys (monkey P, grey circles; monkey R, black squares). **(E)** Same as (B) for the selected series from adaptive decoder dataset. In the adaptive decoder, early (cyan) and late (purple) refer to days, as in Fig. 1C and Fig. 2D. **(F)** Same as (C) for the selected series from adaptive decoder dataset. **(G)** Comparison of 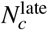and 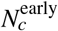 as a fraction of total readouts (late < early, *** *p* = 9.8 *×* 10−4, *n* = 10, one-sided Wilcoxon signed-rank test). Monkey J: black circles; monkey S: grey squares. **(H)** Classification accuracy as a function of readout population size (combinatorial NAC) for early day. Combinations with top 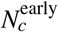 units (deep purple); without top 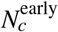 units (orange); with a subset of top 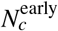 units (grey). Distance between distributions with and without top 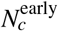 units defined as the discriminability index *d′*. **(I)** Same as (H) for late day. **(J)** Comparison of distance between distributions between early and late learning 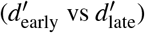 across series (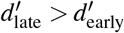, ** *p* = 1.9 ×10−3, *n* = 10, one-sided Wilcoxon signed-rank test). Formatting as in panel G.

We analyzed NACs from longitudinal BCI studies with both fixed [16] and adaptive decoders [28] to assess whether task encoding was concentrated in a few neurons or distributed across the readout population across learning. On each day (early, late), we quantified the classification accuracy of a unit in isolation (Fig. 3B,E), and then constructed a rank-ordered NAC by adding units to our decoding ensemble from most to least predictive (Fig. 3C,F). Since early fixed decoder performance was poor, we grouped data into epochs of 150 trials (early: first 150, late: last 150), consistent with previous analysis [29, 30]. Adaptive decoder datasets allowed analysis across days due to sufficient early trials. Conclusions from our analyses were not significantly impacted by the definitions of early and late datasets.

The relationship between classification accuracy and the number of ranked readout units required for prediction differed between fixed and adaptive decoders. Overall classification accuracy improved between early and late learning (Fig. S3A). We therefore examined the number of units contributing to task encoding by normalizing each day’s NACs (Fig. 3C,F) and then computing the number of neurons (*N*_*c*_) required to reach 80% of the normalized classification accuracy at each learning stage. In fixed decoders, late *N*_*c*_ exceeded early *N*_*c*_, indicating broader distribution of target information with learning (Fig. 3C,D). In contrast, adaptive decoders showed the opposite trend, with a consistently smaller late *N*_*c*_ than early *N*_*c*_ (Fig. 3F,G, Fig. S3E). This effect in adaptive decoder data was even larger when including all recorded units (readouts and nonreadouts, Fig. S3B). We did not observe this effect in arm movement tasks, suggesting that the changes were specific to BCI learning and not due to recording quality (Fig. S3C). Beyond this within-series (early vs. late) trend, *N*_*c*_ qualitatively decreased across chronologically-ordered series for each animal (Fig. S3D). There was no clear relationship between the reduction in *N*_*c*_ and the number of readout units added or removed by decoder perturbations (Fig. S3F), consistent with credit assignment computations occurring on longer timescales rather than tied to any individual decoder intervention.

Next, we examined whether this reduction in *N*_*c*_ carried over to the encoding of kinematic variables in addition to target identity. We found that a smaller subset of readout units, identified using the classification model, was sufficient to accurately reconstruct cursor velocities offline late in learning compared to early (Fig. S3G-I and Supplementary Methods). This suggests that the same units which increasingly contributed to target encoding during learning also contributed to skilled movement. We term this reduction in the number of units needed to accurately decode behavior over time “compaction” to indicate that the brain appears to progressively encode task information in a smaller number of neural units when learning is assisted, in contrast to fixed decoder scenarios (Fig. 3D,G).

Lastly, we asked whether compaction reflected a situation where task information was encoded within a specific subset of readout neurons, or a more general increase in encoding efficiency. We performed a combinatorial variant of the neuron-adding curve analysis, in which we computed classification accuracy for all combinations of units as we vary the number of units used to decode (within readout units only). On each day, we identified the top *N*_*c*_ readout units and labeled combinations that contained all top units (purple) or did not contain any of the top units (orange) (Fig. 3H,I). Early and late in learning, combinations with top units held the most classification power (Fig. 3H,I). Comparing the distributions between early and late revealed that the overall increase in classification power was driven solely by the increase in classification power from top units. We quantified the performance gap between combinations with or without the top 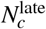 units using a discriminability index (*d*^*′*^) (see Methods). This performance gap increased with learning across series (Fig. 3J), showing that learning-related changes are targeted to a specific subset of readout neurons in BCIs with adaptive decoders.

### Assistive decoder adaptation contributes to learning more compact representations: modeling analysis

Our analysis revealed that neural representations became more compact over multiple days of BCI training with assistive decoder perturbations (Fig. 3E-G), but not without them (Fig. 3B-D). However, experimental BCI data may miss potentially confounding population-level mechanisms due to practical limitations, like sparsely sampled neural activity and an inability to measure *in vivo* synaptic strengths. We therefore simulated BCI task acquisition using an artificial neural network (Fig. 4A) to directly compare learning strategies for identically initialized neural networks trained with fixed or adaptive decoders.

**Figure 4.**
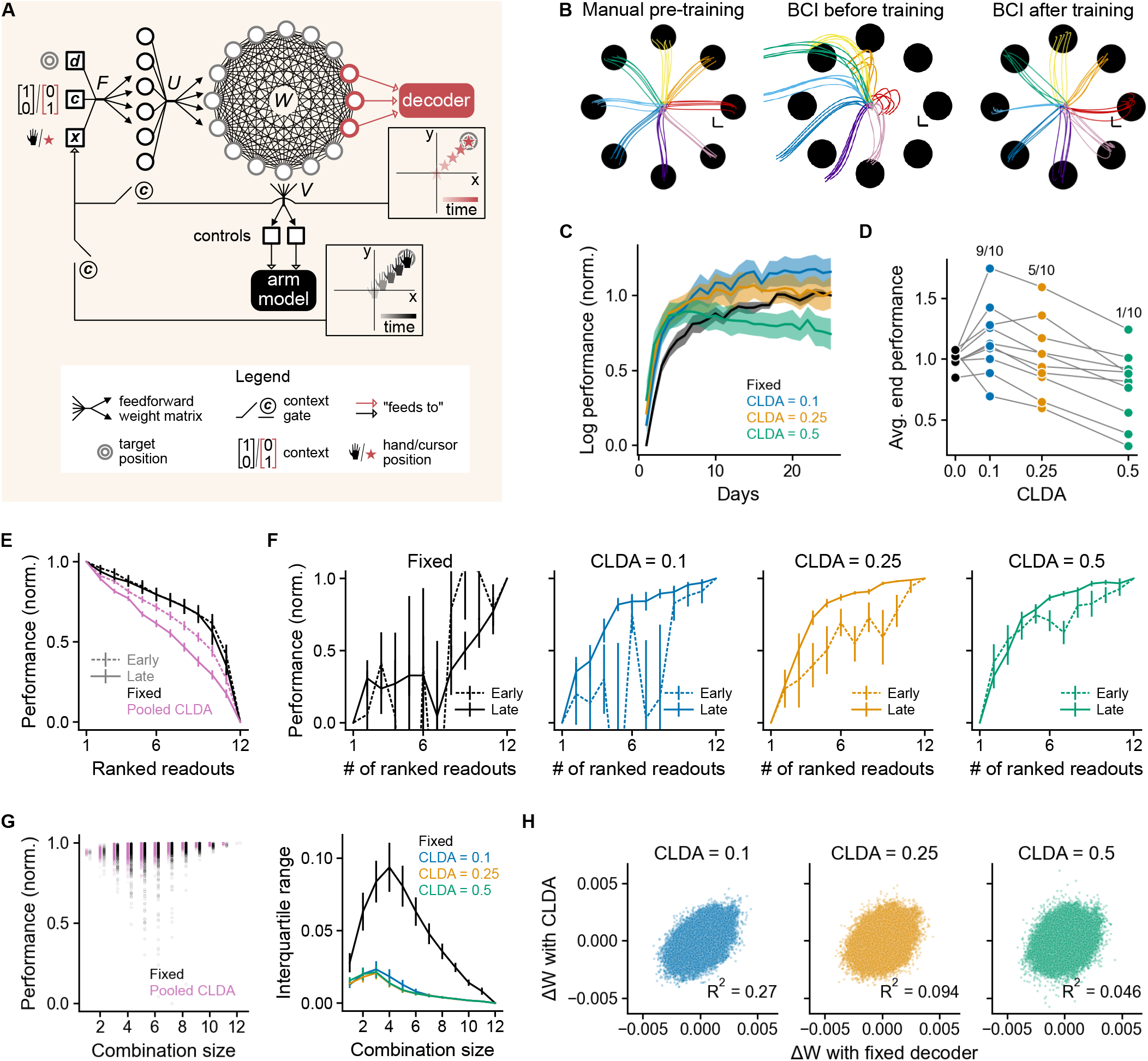
Decoder adaptation contributes to learning compact representations: modeling analysis. **(A)** The RNN received visual feedback about target (**d**) and end-effector (**x**) positions. Based on context (**c**), it controlled either an arm model with all units or a BCI cursor with readout units (red). *F, U, W* and *V* indicate the encoding, input, recurrent and output weight matrices, respectively. See text for details. **(B)** Learning stages. Left: train on arm movement task. Middle: fit decoder parameters to arm trajectories. Right: set context to ‘BCI’ (here CLDA intensity = 0.25) and train on BCI task. Scale bars = 1 cm. **(C)** Normalized log performance (Methods, Eq. 7) for different CLDA intensities. **(D)** Normalized log performance averaged over last 5 days. Ratios at top are the fraction of seeds with better performance with adaptive decoders compared with fixed decoders. **(E)** Ranked single-unit normalized performance (Methods, Eq. 8). Results for adaptive decoders were pooled across CLDA intensities (pink lines). **(F)** Neuron adding curves (NACs) using online performance (Methods, Eq. 9). **(G)** Late combinatorial NAC for a given random initialization of the network parameters (left) and interquartile ranges of the distributions across realizations (right). **(H)** Change in recurrent weights (Δ*W*) with CLDA vs. corresponding change with a fixed decoder. In panels C, E, F and G (right), we plotted ± mean SEM, for *n* = 10 seeds, i.e. random initializations of the network.

We developed a minimal yet biologically plausible RNN model representing a motor cortical circuit that captured the basic learning dynamics of BCI experiments (Fig. 4A). We first trained the network on a center-out arm movement task (Fig. 4B, left) to establish a plausible inductive bias for subsequent BCI training and to mirror experiments [28]. The network received context information about task type (arm vs. BCI) along with the effector and target positions. After initial training, 12 units out of 100 were selected as readouts and a velocity Kalman filter (see Methods) was fitted to manual reach trajectories (Fig. 4B, middle). The context then switched to ‘BCI’ and the network was trained on the BCI task (Fig. 4B, right). In all contexts the REINFORCE algorithm [42] updated recurrent (*W*, Fig. 4A) and input (*U*) weights, while the encoding (*F*) and output (*V*) matrices remained fixed after random initialization. REINFORCE does not rely on backpropagating task errors through the effectors, making it more biologically plausible.

Adaptive decoding in the model followed the principle of CLDA algorithms used in experiments [28], focusing on decoder updates when readout ensemble remained unchanged (see Methods). CLDA was performed on each “day”, corresponding to seeing each target 100 times. Regardless of CLDA intensity (a factor representing how much the decoder changed; Eq. 2), adaptive decoders accelerated early performance improvements compared to fixed decoders (Fig. 4C). Moderate CLDA intensities maintained or improved final BCI performance (Fig. 4D) while preserving generalization properties comparable to fixed-decoder training (Fig. S4A). However, excessive adaptation degraded final performance (Fig. 4D, CLDA = 0.5), likely because CLDA’s objective (reach towards the target) conflicted with the task objective (reach the target with vanishing velocity in a specified time) late in training (Methods, Eqs. 4-5). Stopping high-intensity CLDA once a performance threshold was reached mitigated this issue (Fig. S4B). Overall, in the model, decoder adaptation facilitated rapid BCI task acquisition, as in experiments. Our results also underscore potential shortcomings of long-term CLDA applications at high intensity.

Next, we explored whether adaptive-decoder learning led to more compact representations in the model. While this was assessed offline in the data (Fig. 3), the model allowed for an online evaluation using the task loss as the performance metric (Methods, Eqs. 4-5). Each individual readout unit controlled the cursor in turn and its reach performance was evaluated both ‘early’ (end of first day) and ‘late’ (end of last day). Readout units were then ranked according to their performance. Early and late ranked single-unit performances (Methods, Eq. 8) differed only for adaptive decoders (Fig. 4E and Fig. S4C,D), with faster decrease late in training compared to early (Fig. 4E, pink lines). Thus, the dominant units became relatively more dominant with learning on average under network-decoder co-adaptation. Using these rankings, we then computed a ranked NAC (Methods, Eq. 9). Consistent with our experimental findings, adaptive decoders contributed to the compaction of representations of task performance (Fig. 4F and Fig. S4E) and kinematic variables (Fig. S4F). With a fixed decoder, ranked NACs were highly variable across seeds, with no sign of compaction as 11 (out of 12) ranked readout units were required to reach 80% of the maximal performance late in training (Fig. 4F, left). Notably, stopping CLDA early not only preserved performances (as mentioned above), but also tended to prevent the emergence of compact representations (Fig. S4G). Overall, our model confirms and expands on our observations that decoder adaptation singles out certain units for BCI control and progressively shapes more compact neural representations.

Using the online loss metric, we next obtained combinatorial unit-adding curves (Fig. 4G, left) and computed the corresponding interquartile ranges (IQR; Fig. 4G, right). These IQRs measured the dispersion of performance for each combination size (Methods, Eq. 10). The IQRs were significantly greater for a fixed decoder, which was caused by few units with very low contributions to the overall performance (Fig. 4E and Fig. S4C). Therefore, while decoder adaptation promoted more compact representations with a few dominant units, it also mitigated the impact of unreliable ones.

The model suggests that neural plasticity alone, without decoder adaptation, does not lead to compaction. Thus, decoder adaptation must interact with the model’s plasticity to generate this feature of neural representations. To show this, we calculated the total changes in recurrent weights across learning, Δ*W*, with and without decoder adaptation, and computed their coefficients of determination (Fig. 4H). All connections—among and across readout and nonreadout units—were included. Since our simulations were precisely matched in terms of random number generation across CLDA intensities, comparing these weight changes one-to-one was meaningful. The coefficients of determination decreased as CLDA intensity increased, indicating that the total weight changes with an adaptive decoder became increasingly uncorrelated from those obtained with a fixed decoder. These results suggest that brain-decoder co-adaptation drives distinct neural plasticity patterns and contributes to compaction.

### Compact representations encode task information in low-variance population modes

Our neural data analysis and model revealed that compaction at the unit level in neural representations emerge as the result of long-term learning processes occurring alongside an adaptive decoder. Does this reshaping of sensorimotor encoding at the level of individual neurons reflect a broader reorganization within the neural population? Many studies represent coordinated neural activity with principal components (PCs) capturing the most prominent directions of variability [43, 44, 45]. In this view, each PC can be treated as an independent component or “mode” of neural population activity.

We explored how compaction changed population structure by performing a variation of our neuron-adding curve analysis in which population-level modes served as features for classification rather than individual units (Fig. 5A). For early and late days, we performed principal component analysis, ranked PCs based on their classification accuracy (Fig. 5B) and computed a rank-ordered PC adding curve (Fig. 5C). Similar to the compaction observed at the unit level, we found that the number of PCs required for 80% classification accuracy decreased with learning (Fig. 5C,D). This observation is not a trivial consequence of unit-level compaction (see Supplementary Discussion), and confirms compaction of representations at the level of populations.

**Figure 5.**
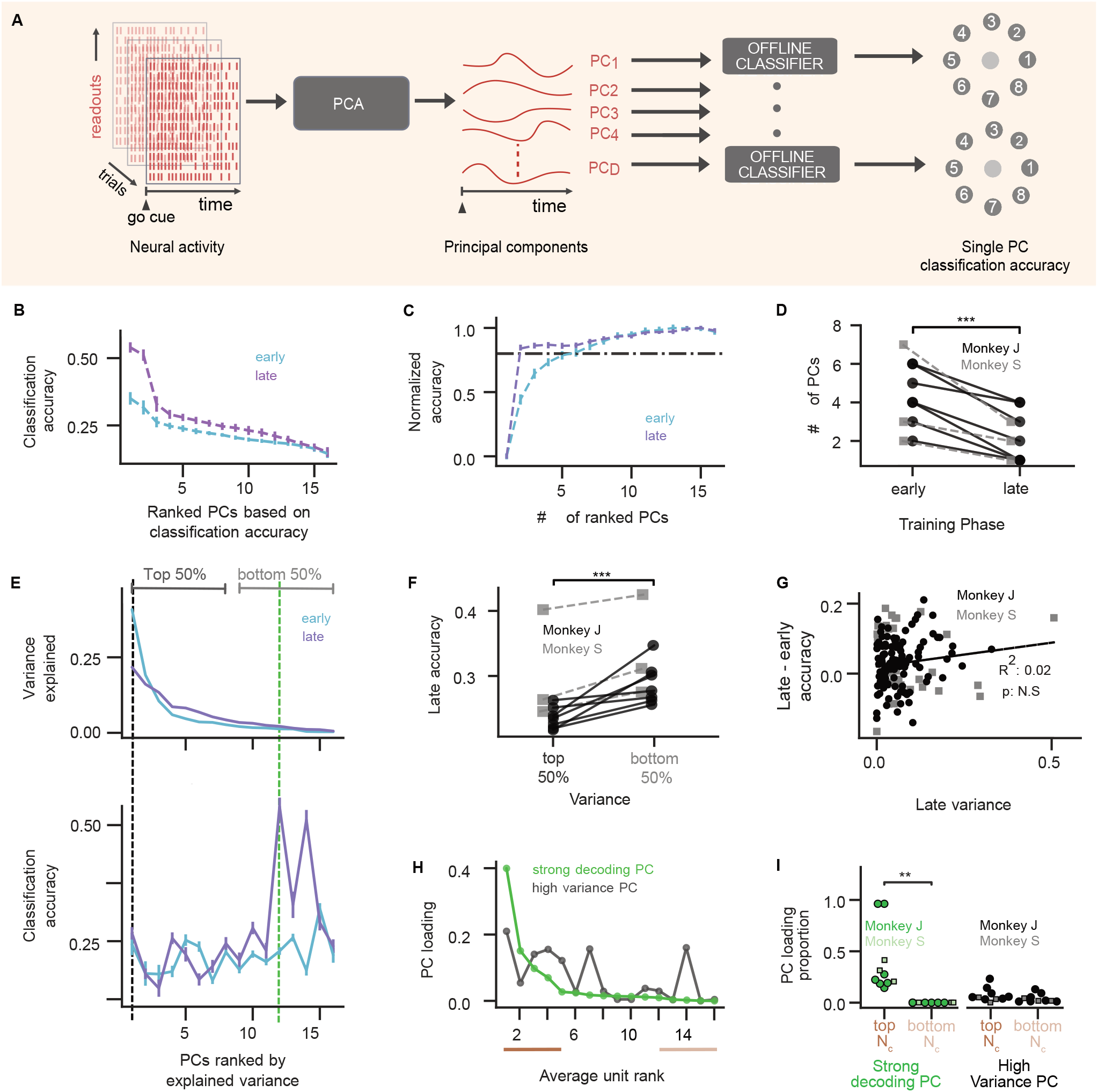
Task information emerges in low-variance modes. **(A)** Decoding analysis with PC modes (see text). **(B)** Individual PC classification accuracy (similar to Fig. 3E) for early (cyan) and late (purple) learning phases from selected series. PCs ranked by classification accuracy. **(C)** Ranked dimension adding curve (similar to ranked NAC in Fig. 3F) for early and late days for the example series. **(D)** Comparison of number of PCs needed for 80% classification accuracy on early and late days across series (*** *p* = 9.7 ×10^−4^, *n* = 10, late *<* early, one-sided Wilcoxon signed-rank test). Monkey J: black circles; monkey S: grey squares. **(E)** Top: Variance explained by each PC (ranked by variance explained). Bottom: Classification accuracy per PC, ordered by variance explained, Error bars denote 95% confidence intervals from 104 random draws of trials. The strongly decoding PC (green) has the highest accuracy. **(F)** Comparison of mean classification accuracy from top 50% vs bottom 50% PCs during late learning day across series. Top 50% is lower than bottom 50%: ** *p* = 0.001, *n* = 10, one-sided Wilcoxon signed-rank test. Data formatting as in D. **(G)** Change in classification accuracy (late − early) plotted versus late variance per PC. No significant trend: *R*2 = 0.01, *p* = 0.26, *n* = 10. Data formatting as in D. **(H)** Square of PC loadings from each readout unit for the strong decoding PC (green) and high variance PC (black). Units on the x-axis are ranked by their average classification performance. **(I)** Comparison of sum of squares of PC loadings from top 4 ranked readouts and bottom 4 ranked readouts from strong decoding PC and high variance PC across all series. Strong decoding PC: ** *p* = 0.002; high variance PC: *p* = 0.38 (ns), two-sided Wilcoxon signed-rank test, *n* = 10.

We further observed that variance became more broadly distributed across PCs in the readout population after learning, suggesting an increase in dimensionality (Fig. 5E, top). To validate this observation, we calculated the dimensionality of readout activity using the participation ratio, a covariance-based dimensionality measure [46, 47], normalized by the number of readouts (PR_norm_). Unlike in natural motor learning studies where neural variability tends to shrink [31] or fixed-decoder BCI studies where “shared” dimensionality decreases [29, 30], dimensionality remained the same or increased during BCI learning with adaptive decoders (Fig. S5A,C). Re-analyzing fixed-decoder data using PR_norm_ confirmed that dimensionality changes with learning differ in the presence of adaptive decoders (Fig. S5B,C).

How can neural representations become more compact while dimensionality does not decrease? We linked variance captured by population-level modes with task information across learning. While variance was more distributed across modes after learning (Fig. 5E, top), modes capturing the highest population variance were not necessarily the most predictive (Fig. 5E, bottom). We quantified the relationship between variance explained and task information for population modes by dividing PCs into two groups for each learning series: the top 50%, which explained approximately 80-84% of population variance, and the bottom 50%, which explained 16-22% of population variance. Across all learning series, the average classification accuracy on late days was higher in the bottom 50% group compared to the top 50% group (Fig. 5F), showing that task information was more often found in low-variance modes after learning. Moreover, the changes in a PC’s classification accuracy during learning were not correlated with its variance explained (Fig. 5G). Thus, compaction co-occurred with a nondecreasing dimensionality because task-relevant information became embedded in the low-variance modes.

Lastly, we examined whether, after learning, task-predictive units contributed preferentially to task-predictive population modes. We ranked readouts based on their single-unit predictive power and then quantified their influence on PC modes using the squared PC loadings (see Methods). Comparing PC loadings for an example high-variance PC with those of a strongly decoding PC revealed starkly different patterns of unit contributions (Fig. 5H). As expected, strongly decoding PCs receive large contributions from strongly decoding units. In contrast, high-variance PCs have contributions from a seemingly random mix of units. To quantify this effect across all learning series, we compared the sum of squared PC loadings from the top *N*_*c*_ task-predictive units to the bottom *N*_*c*_ task-predictive units for the strongest decoding PC and highest variance PC in each series (Fig. 5I). Strongly decoding PCs consistently drew more from the top-performing units, with negligible input from the least predictive units, a pattern not observed in high-variance PCs (Fig. 5I). Thus, the most predictive PCs in neural representations learned with adaptive decoders were not only associated with moderate to low variance but also composed primarily of the most task-predictive units. This indicates that task-relevant information can be compactly encoded in low-variance population modes through the coordinated contributions of highly informative units, challenging conventional views on the relationship between neural variability and task encoding.

## Discussion

We found that assistive BCI perturbations led to a compaction of neural representations, concentrating task information into fewer neurons and neural modes that capture a relatively small portion of the total population variance. Both experimental data and our model suggest that compaction does not occur without these assistive perturbations. By revealing new impacts of BCI *decoders* on the brain’s *encoder*, our work also opens new opportunities to design BCI algorithms that deliberately steer a two-learner system toward better task representations.

Our findings highlight the complexity of “error” signals in BCI learning. Past BCI studies emphasized links between neural changes and task performance. Neural activity may only reorganize when decoder perturbations introduce task errors and violate existing neural correlations [15, 48]; and adapting a decoder to decrease task errors reduces neural changes over time [28]. Unlike most prior studies [14, 15, 35, 48, 49], here the decoder changes did not introduce overt *task* errors, instead creating intermittent changes to the mapping between neural activity and behavior that do not always have an impact on task performance. Adaptive decoders, therefore, manipulate the mapping between *intent* and *control*, even when task error is not affected. We found that these decoder changes impacted learned neural representations (experiment, Fig. 3B-G and model, Fig. 4E-G) and synaptic-level changes (model; Fig. 4H). This is consistent with our hypothesis that decoder adaptation interacts with error-driven skill learning computations, and suggests that learned representations in BCI tasks are influenced by error signals beyond that of task objectives. Motor learning studies suggest that skill acquisition involves both fast (within-session) and slow (multi-day) adaptation [1, 2, 35, 50] with different error sensitivities [51]. Fast and slow adaptation likely involve different neural changes, such as fast adaptation via minimal synaptic changes [49, 52]. One possibility that can be explored in future studies is that the initial performance boost from adaptive decoders could supplant rapid learning mechanisms driven by task errors, while slow mechanisms that shape synaptic changes remain and are impacted by the algorithm. This is consistent with our observation that compaction emerged over time, rather than explicitly tied to individual decoder update events (Fig. S2H, S3F), and trends of more compact representations *across* learning series (Fig. S3D). Indeed, adaptive decoders may alter slower credit-assignment processes using error signals generated through mechanisms like an “internal model” [53, 54].

Our results also indicate that credit-assignment computations during extended BCI practice operate at scales spanning single neurons as well as coordinated neural populations, and that these computations are impacted by adaptive decoders. Importantly, compaction of population modes and that of units do not trivially follow one another in general (see also Supplementary Discussion), making it noteworthy that dominant units preferentially contribute to strongly-decoding modes (Fig. 5H-I). Our offline classification of target identity with neural data (Fig. 3, Fig. 5) and online assessment of neuronal contributions to control in the model (Fig. 4) revealed that adaptive decoders strongly influence which neurons or neural modes participate in the task, leading to exaggerated credit assignment. These task informative neurons appear to form an exclusive coordinated population, since task informative population modes have minimal contributions from other neurons. This exclusion, in turn, partially explains why modes with task information ultimately capture relatively little of the overall population variance. This is consistent with our supplemental analysis comparing variance and dimensionality changes during learning between fixed and adaptive decoders. Neural variability and dimensionality have been found to reduce with practice (along with behavioral variability) in both motor learning [31] and BCI learning with fixed decoders [29, 30]. We found that this is not the case when the decoder adapts, instead leaving population dimensionality and variability largely unchanged (Fig. S5BC). Together, these results suggest that assistive decoders may shape neuron-level learning processes in a way that helps maintain population dimensionality. Our study adds to observations that neural variance and task information are not necessarily linked [55], showing that long-term learning phenomena can occur in relatively low-variance modes where they may easily be missed by linear-based variance analyses [40].

Our study reveals new challenges and opportunities in designing long-term BCIs. Given measurement instabilities over a BCI’s lifetime, some form of adaptive decoding is unavoidable. Our results highlight that long-term learning mechanisms will likely occur alongside decoder changes, making long-term BCI systems inherently *co-adaptive* [25, 56]. Moreover, our results demonstrate the inherent entangling of *decoding* and *encoding* in co-adaptive systems. The adaptive decoding algorithms used in our experiments [20]—closely related to widely used adaptive algorithms [11, 21, 23, 24, 26, 36, 37]—produced compaction of neural representations. This suggests that the brain-decoder system ultimately relies on a smaller set of neural features, which will impact BCI performance. For instance, compact representations may improve computational efficiency, and could lead to task encodings that are more easily separated from other behaviors to reduce interference from other task demands [28]. Compact encoder-decoder systems may also protect against unreliable units (Fig. 4G). Yet, the dominance of a few features can prevent more subtle information from influencing learning [57] and compact task encodings may increase sensitivity to neural recording loss. Our model highlights the opportunity to balance these potential trade-offs by properly fine-tuning the decoder adaptation algorithm. Such models will likely be valuable for more principled designs of co-adaptive BCIs [56]. Understanding the mechanisms by which assistive decoders shape task encoding and their functional implications will ultimately make it possible to design BCIs that provide high performance for someone’s lifetime.

## Acknowledgements

The authors thank Jose M. Carmena, who shared data collected in his laboratory for this study, and Vivek Athalye for his assistance with the fixed decoder dataset. The authors’ research was supported in part by an IVADO postdoctoral fellowship, Canada First Research Excellence Fund/Apogée (AP), an NSF Accelnet INBIC fellowship (PR), a Simons Collaboration for the Global Brain Pilot award (898220, GL and ALO), a Google Faculty Award (ALO and GL), an NSERC Discovery Grant (RGPIN-2018-04821, GL), the Eunice Kennedy Shriver National Institute of Child Health and Human Development (NIH K12HD073945, ALO), and NIH grant R01 NS134634 (ALO). GL further acknowledges support from the Canada CIFAR AI Chair Program and the Canada Research Chair in Neural Computations and Interfacing (CIHR, tier 2). The content of the present paper is solely the responsibility of the authors and does not necessarily represent the official views of the funding agencies.

## Methods

### Experiment

#### Neural data and behavioral task

We analyzed neural activity as two male rhesus macaques (*Macaca mulatta*, monkeys S and J) learned to move a 2D cursor controlled via neural activity to perform a point-to-point reaching task while the relationship between neural activity and cursor movement was intermittently adapted across days (Fig.1C). The task required animals to initiate a trial by moving the cursor to the central target. After reaching the center, one of eight peripheral targets appeared. After a brief hold at the center, a target color change cued the animal to start reaching. Successfully completing a trial required the monkey to enter the peripheral target within a specified reach time (3 − 10 s) and hold the cursor inside the target briefly (250 − 400 ms). The monkey then received a liquid reward. Because we were primarily interested in neural representations of movement direction, we analyzed all trials in which the animal successfully reached the peripheral target regardless of whether the peripheral hold was successful or not.

Both monkeys were implanted with 128 microwire electrode arrays in motor and premotor cortices to record spiking activity. Spike-sorted multi-unit activity (monkey S) or unsorted threshold crossings defining channel-level multi-unit activity (monkey J) were used for BCI control. We refer to neural activity for both animals as “units”. A subset of recorded units was used for the real-time BCI cursor control, which defines the readout neural population. Readout neurons were chosen based primarily on their stability across days, assessed qualitatively by experimenters by observing recordings over time. Functional properties, such as tuning for arm movements or offline task predictive power, were not used to inform readout unit selection. Units that were recorded on the same array, but not used as inputs to the BCI decoder for cursor control, define the nonreadout neural population. Stable readout units were defined as units consistently used throughout an entire BCI series.

All procedures were conducted in compliance with the NIH Guide for the Care and Use of Laboratory Animals and were approved by the University of California at Berkeley Institutional Animal Care and Use Committee.

#### BCI control with adaptive decoders

Readout neural activity controlled the 2D cursor using a position-velocity Kalman filter (KF) [20, 23, 28, 58]. KF parameters were trained and intermittently updated using previously-developed algorithms that estimate parameters during closed-loop BCI control (closed-loop decoder adaptation, CLDA) [20]. CLDA re-estimates the parameters of the KF using the subject’s neural activity and estimated motor intent during BCI control and updates the decoder used for BCI control in real-time. Updates to the KF BCI used the SmoothBatch algorithm [20] (monkey J; see below) which updates a subset of the KF parameters or the ReFIT algorithm [23] (monkey S) which updates all KF parameters.

Monkeys practiced with a given decoder mapping (and subsequently updated versions of that mapping) for multiple days, defining a learning series. CLDA served two main purposes: (1) to enhance closed-loop performance from the initial decoder, and (2) to maintain performance despite shifts in neural activity, such as the loss of a unit within the BCI readout ensemble. The initial CLDA session (day 1) of a decoder series lasted approximately 5–15 minutes, providing the subject with sufficient performance to reach all targets successfully. Intermittent updates included (1) alterations to decoder parameters without changing the readout unit identities (weight change only) if there was a noticeable performance drop, assessed as a success rate decrease of approximately 10%–20%, and (2) interventions where readout unit identities were altered to address loss of a unit previously in the readout ensemble along with decoder weight changes (readout + weight change) (Fig. S1A). In these cases, CLDA was briefly run for 3–5 minutes, corresponding to 1-2 updates of the KF weights. Readout unit swaps and weight adjustments were made only if previously used readout units could no longer be isolated or identified. When a performance drop from the previous day was observed but all readout units remained present, only weight changes were made without altering readout units. CLDA therefore had two primary effects: initial CLDA led to large decoder changes that boosted performance to facilitate reaches to all targets, akin to initial “calibration”; mid-series CLDA made notably smaller adjustments to decoder parameters, which primarily maintained or modestly improved performance [28]

Monkey J and monkey S learned 13 and 6 adaptive decoder mappings in total, respectively [28]. Our analysis focused on extended learning series (longer than four days) that included intermittent CLDA on the KF decoder (monkey J: 7, monkey S: 3). Each learning series varied in length and continued until performance plateaued. Longer series tend to have more CLDA events (Fig. S1D). Information about the number of readouts and CLDA events in each learning series is shown in Supplementary table 1. Monkey S had a reduced frequency of mid-series CLDA, which does not necessarily imply better task performance than Monkey J, but may potentially reflect more stable neural recordings. Several of Monkey S’s series were shorter than Monkey J’s, which may have contributed to the reduced number of CLDA events. The original study intentionally manipulated the degree of initial CLDA to vary BCI performance on Day 1, with each learning series having different initial performances based on the extent of decoder adaptation (Fig. 1C, Fig. S1C). Some series began with lower initial performance due to minimal or no CLDA applied at the outset. Full details on adaptive decoder methods and the dataset are provided in Orsborn *et al*. [28].

### Data analysis

#### Learning analyses and behavioral performance metrics

We quantified long-term learning changes during a series by comparing “early” and “late” training days. We defined an early training day as the first day with at least 25 trials per target direction. A late training day was the last day in the learning series with at least 25 trials per target direction.

We assessed behavioral improvements using standard performance metrics such as mean success percentage and reach times (Fig 1C). Success percentage was computed as the fraction of rewarded trials to the total number of trials initiated. Reach time was the time between cursor leaving center target and entering peripheral target.

#### Neural data preprocessing

Neural activity was binned at 100 ms and trial-aligned to the go-cue. We included neural activity from the onset of the go-cue to 1800 ms after as inputs to an offline decoding model (see below). Altering the time included (from go-cue to a range of 500 to 1800 ms post go-cue) was not found to qualitatively change reported findings (not shown). Because reach times vary over the course of learning, trials with a post-go-cue duration shorter than 1800 ms were zero-padded; this was most common in late phases of learning.

#### Logistic regression model

We analyzed changes in neural representations with multiclass logistic regression (LR) models that used neural activity (Figs. 2, 3) or transformations thereof (Fig. 5; see below), to predict target identity. The classifier model received flattened activity vectors composed of the activities of all units across time bins between go-cue to reach completion for each trial, and generated an output corresponding to the predicted target label. To control for variability in the number of trials completed to each target across days, we generated a training set where trials were randomly selected (without replacement) to match the number of trials to each target across days. We selected 25 trials per reach direction (200 trials per day), matching our criteria for defining “early” training phases. The remaining trials on a day were used as a test set. Unit firing rates were z-scored across trials for each set separately. We performed 100 training-test splits on each day, randomly selecting 200 trials for training (25 trials per reach direction) and designating the remaining trials for the test set in each draw. This approach allowed us to calculate the 95% confidence interval on classification accuracy (displayed as error bars). We used the mean accuracy across the training-test splits to compare classification accuracy across series and animals (Fig. 2E).

We implemented our LR models using the Python library Scikit-learn [59] with the L-BGFS solver and L2 regularization (hyperparameters: C = 1, max_iters = 1000). We did not observe any considerable changes in classification accuracy between LR models with or without regularization on late days (Fig. S2A). The hyperparameter C represents the inverse of regularization strength, with smaller values indicating stronger regularization, which helps prevent overfitting by penalizing large coefficients. We conducted a grid search for C ranging from C = 1e-5 to 1, across different neural populations (all, readouts, and nonreadouts). The purpose of this analysis was to confirm that our results were not merely due to the model performing unevenly across these groups. We found that classification accuracy was not more or less sensitive to the choice of C for any particular neural population (Fig. S2B). We note that this is not an analysis aiming to optimize the model’s classification performance for each population.

Parameter values (model weights) for a single training-test split and for a single target (target 2) were used to illustrate the LR model (Fig. 2B). We quantified the similarity of decoder weights across days by calculating the similarity between the model weight matrices *W* on consecutive days, *W*^*d*^, *W*^*d*−1^ (Fig. 2C), similar to previous analyses [16, 28]. This evaluation was performed by calculating the correlation coefficient between the flattened matrices of model weights (weights for each target concatenated) for each day.

#### Rank-ordered neuron adding curve

We estimated the contribution of each individual unit to target identity prediction by running a LR model for each unit separately (Fig. 3A, left). We then ranked units based on their classification accuracy (Fig. 3A, middle and Fig. 3B). A ranked-ordered Neuron Adding Curve (NAC) was produced by calculating the classification accuracy with the highest ranking unit, then with the first and second highest ranking units, and so on (Fig.3A, right and Fig.3C,F) [41]. These analyses used the same training-test split protocol described above. To account for the lower maximum classification accuracy on early days compared to later days, we normalized the NAC classification scores to each day’s maximum performance using a linear scaling. We then computed the number of ranked units (*N*_*c*_) required to achieve 80% normalized prediction accuracy for early and late days. We performed a Wilcoxon signed-rank test using the *N*_c_ ‘s across series and monkey with alternative hypothesis 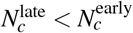 (Fig. 3D,G).

The “combinatorial” NAC, as described in the main text, tested all possible combinations of units for each size of ensemble. We evaluated the separability between the classification accuracy distributions with and without the top *N*_*c*_ units (Fig. 3H-J) using a discriminability index, *d*^*′*^ = |*m*_1_ −*m*_2_|*/*[(*s*_1_ + *s*_2_)*/*2], where *m*_*i*_ and *s*_*i*_ are the mean and standard deviation of distribution *i*, respectively. A higher *d*^*′*^ indicates that two distributions are more separable.

#### Classification using neural modes

We performed principal component analysis (PCA) on the readout activity for the early and late day separately (Fig. 5A). All trials with completed reaches were used for PCA each day, and readout firing rates were z-scored per day. We constructed PC adding curves and analyzed compaction in the PCs in the same way as in the above rank-ordered NAC, but using PCs instead single-unit activity (Fig. 5B-D). Importantly, we kept all PCs (no dimensionality reduction) and thus the number of PCs was equal to the number of readouts. We compared classification accuracy in low and high-variance PCs by splitting PCs into two groups (top 50% and bottom 50%) (Fig. 5F). These groups were defined by ranking PCs according to variance explained and assigning the first (last) *N/*2 PCs into the top (bottom) group.

#### PC loadings

The loadings are the components of the vectors produced by PCA, which generate the PCs by projecting a random vector variable onto them [60]. After normalizing the loading vector, we took the square of a readout’s loading to represent its relative contribution to the PC. For each learning series, on late days, we identified the PC with the highest variance and the PC with the strongest classification accuracy (Fig. 5E), and the corresponding loading vector. For these PCs, we plotted the squared PC loading for each unit against its average rank based on its classification accuracy (Fig. 5H). We calculated the average rank for a unit from the distribution of their ranks across the training/test splits. Finally, we computed the sum of squared loadings (that we called the PC loading proportion) for the *N*_*c*_ leading and trailing units, according to their average rank (Fig. 5I).

### Statistics

To evaluate statistical differences between early and late phases of learning, we applied the Wilcoxon signed-rank test for pairwise comparison. Significance levels are indicated as follows: ****: *p* ≤ 10^−4^, ***: 10^−4^ *< p* ≤ 10^−3^, **: 10^−3^ *< p* ≤ 10^−2^, *: 10^−2^ *< p* ≤ 0.05, ns: *p >* 0.05. The absence of a statistics comparison line in figures between two groups indicates that no statistically significant difference was found. Unless specified otherwise, the sample size for all such early vs late comparisons was *N* = 10 learning series for the BCI task, consisting of 3 series from monkey S and 7 series from monkey J. Inclusion of all learning series as opposed to only extended (*>* 4 days) series, did not qualitatively change the trends observed (not shown). In the supplementary material, control data from *N* = 6 learning series obtained from monkey J related to the arm movement task were used (Suppl. Figs. S2D,E and S3C).

### Model

Simulations were implemented in C++ using the Eigen library [61]. Analyses were performed in Python, using the Numpy and Scipy packages. Values for parameters are included in Table 1. Code will be made publicly available upon publication.

#### Arm model

Our RNN model was initially trained to control a planar torque-based arm [62] consisting of a shoulder and an elbow joint. The arm only moved in the horizontal *x*−*y* plane. The dynamical variables were the angle of each joint (**q** = [*q*_1_, *q*_2_]^T^), their velocity 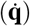 and the applied torque (***τ***). These were linked through the dynamics

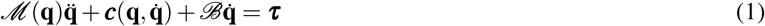

where

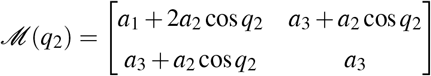

is a 2 *×* 2 inertia matrix,

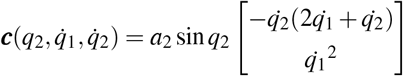

represents the centripetal and Coriolis force vector and

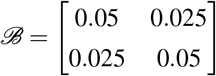

is the joint friction matrix. Parameters *a*_1_, *a*_2_ and *a*_3_ are functions of the moments of inertia (*I*), masses (*m*), lengths (*l*) and distances from joint center to center of mass (*d*) of the two links (see Table 1), with 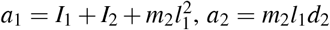 and *a*_3_ = *I*_2_. The torques corresponded roughly to muscle activations and were related to the control signals **u** generated by the network dynamics by 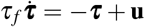 [63]. In simulations, these equations were discretized in time using the standard Euler scheme. The RNN defined below generated controls **u**_0_, …, **u**_*T*−1_ and the arm’s dynamics produced a sequence of states starting from an initial condition 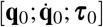 The position, velocity and acceleration of the end effector (hand) are denoted by **x**, 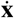 and 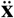, respectively. The initial condition was that the end effector be motionless 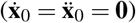 with its initial position **x**_0_ drawn from a uniform distribution centered on the center of the workspace, with radius 0.5 cm. The end effector position was obtained by a transformation of the joint coordinates, with the length of each link, *l*_1_ and *l*_2_, as parameters: **x** = [*l*_1_ cos *q*_1_ + *l*_2_ cos(*q*_1_ + *q*_2_), *l*_1_ sin *q*_1_ + *l*_2_ sin(*q*_1_ + *q*_2_)]^T^. The corresponding end effector velocity was 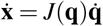, where *J*(**q**) is the Jacobian of the transformation from **q** to **x**. An expression for the end effector acceleration, needed in the learning objective described below, was obtained by differentiating 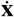 with respect to time.

#### Decoder

The RNN model was then trained to perform BCI control. As in Ref. [28], the BCI decoder was a Kalman filter, using arm trajectories and neural activities to encode the Kalman model parameters, and then inverting the state and observation equations (defined below) to decode newly recorded neural activity. In discrete time, the state and observation equations read

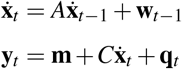

where **q**_*t*_ ∼ 𝒩 (**0**, *Q*) and **w**_*t*_ ∼ 𝒩 (**0**,*W*) are independent Gaussian random vectors, **y**_*t*_ is the firing activity and **m** is the average rate. The dimension of vectors **y**_*t*_, **m** and **q**_*t*_ was the same as the number of readouts in the model, *N*_readout_ = 12. Note that only the end effector velocity 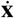 appears here, corresponding to a scenario where uncertainty on velocity does not propagate to position [23]. The position was determined by integrating the velocity.

Matrices *A, W, C* and *Q* were determined from simulation data during arm control following the algorithm described in Ref. [20]. To decode the cursor velocity while measuring neural activity online, we used the update equations

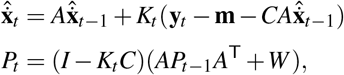

where 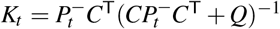 is the Kalman gain with 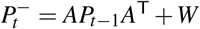. For simplicity, we set the initial estimate 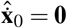 with perfect precision, so that the error covariance matrix was *P*_0_ = 0.

Closed-loop decoder adaptation (CLDA) was implemented following SmoothBatch [20, 28] using data recorded under brain control. Recorded velocities were rotated towards the current target, representing intended velocities [23]. This data with the rotated velocities produced new estimates for matrices *C* and *Q* and vector **m**, denoted Ĉ, 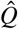 and 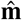, and the values from the last step were replaced according to

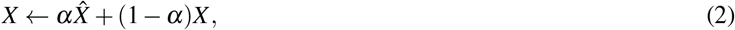

where *X* ∈ {*C, Q*, **m**} and 0 ≤ *α* ≤ 1 is the *CLDA intensity*; *α* = 0 corresponds to a fixed decoder. In the model, CLDA was performed each day irrespective of performance, unlike in experiments, where within-series CLDA was applied only when performance declined.

#### Network model

The RNN contained *N* units fully connected by a recurrent weight matrix *W* (see Table 1). The units’ output were their firing rates **r**, which were obtained from their “membrane potential” **v** by applying an activation function ***φ***(·) elementwise: **r**_*t*_ = ***φ***(**v**_*t*_), i.e., *r*_*t,i*_ = *φ*(*v*_*t,i*_) for each unit *i*. The transfer function *φ* was the rectified linear unit (ReLU), producing non-negative firing rates. The membrane potentials obeyed

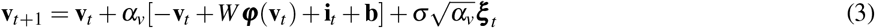

where *t* = 0, 1, …, *T* − 1. These dynamics integrate with time constant 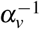 a recurrent input from other units in the network (*W* ***φ***(**v**)), an external input representing information about the task (**i**)—e.g., the position of the target to reach; see below—, a bias (**b**) and a zero-mean Gaussian white noise 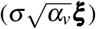. The initial condition for the membrane potential was drawn from a uniform distribution 𝒰 (−1, 1).

The input **i** was given by 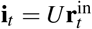, where *U* is the input weight matrix and 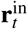 is the activity of the input layer. This input layer activity encoded premotor information. The encoding was a simple random projection of delayed information about target position **d**, end effector position **x** and context **c**—either [1, 0]^T^ for arm control or [0, 1]^T^ for BCI control—followed by the ReLU operation. Symbolically, 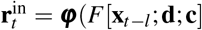), where *F* is a non-learnable random matrix and *l* = 10 is the delay. The output of the network—the controls applied to the arm models—was a linear mapping of the network activity [65]: **u**_*t*_ = *V* **r**_*t*_ = *V* ***φ***(**v**_*t*_), where *V* is the output matrix.

#### Training objective

Let **d**_*k*_, *k* = 0, …, *K* − 1, represent the position of the *K* = 8 peripheral targets at a distance *D* = 7 cm from the center target, with **d**_*k*_ = *D*[cos(2*πk/K*), sin(2*πk/K*)]^T^. The training objective was to minimize

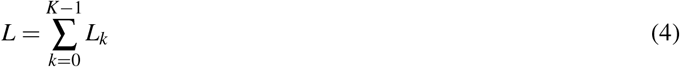

where

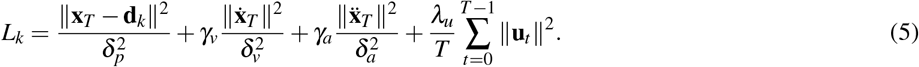

Therefore, the objective was to reach the target at the end of the trial (first term) with vanishing velocity (second term) and acceleration (third term), and with an effort penalty (fourth term). Parameters *δ*_*p*_, *δ*_*v*_ and *δ*_*a*_ were used to rescale the position, velocity and acceleration terms. Hyperparameters *γ*_*v*_, *γ*_*a*_ and *λ*_*u*_ controlled the relative weight of the velocity, acceleration and effort costs with respect to the position loss. Hyperparameters *λ*_*u*_ and *γ*_*a*_ were nonzero only under arm control (see Table 1).

#### Learning

The recurrent weights *W*, input weights *U* and biases **b** were plastic while the output matrix *V* and the encoding matrix *F* were fixed. Learning was performed *via* node perturbation using the REINFORCE algorithm [42, 66]. The noise ***ξ*** independently applied to all units in Eq. 3 evoked small end effector jitters. In reinforcement learning, jitters that increase reward should be reproduced, and network parameters should be updated accordingly. Here, the reward was taken to be minus the learning objective, *R* = −*L*. The gradient with respect to *W* was estimated using

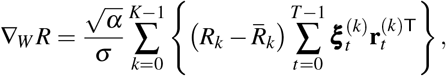

where 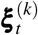 and 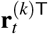 are the noise and activity when target *k* is presented. The reward trace 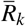 provided an estimate of the expected reward for target *k* [67] and was computed as a moving average

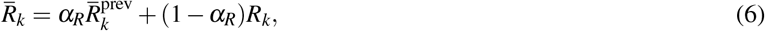

where *α*_*R*_ ∈ [0, 1] is a factor that weights the relative contribution of the present reward (*R*_*k*_) and the expected reward in memory 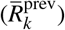 in computing the new reward estimate. For *U*, one simply replace **r**_*t*_ by 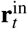; for **b**, the summation is over ***ξ***_*t*_ only. Parameter updates were computed after seeing all *K* targets (an epoch) using Adam updates with standard hyperparameters [68].

#### Analysis of the model

To allow clear comparison across seeds (i.e., random initializations of the network’s parameters) and CLDA intensities, we normalized the BCI training loss *L* relative to the loss with a fixed decoder, denoted *L*^(0)^. For each seed, we computed the log_10_ of *L* and *L*^(0)^ for the last epoch of each day and we defined a new performance metric (“normalized log performance”, Fig. 5C) as

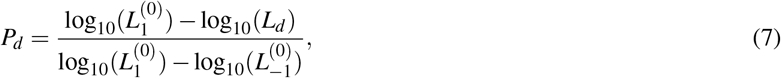

where 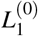and 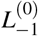 are the losses with a fixed decoder on the first and last day, respectively. This linear transformation sought to transform the loss for the fixed decoder to the interval [0, 1].

Single-unit normalized performances (Fig. 5E) were computed by first evaluating the reach performance for each individual unit and ranking them from the most important to the least important. The performance of each unit was measured by restricting the Kalman filter parameters **m**, *C* and *Q* to that unit and recomputing the Kalman gain *K*_*t*_ accordingly. This ensured that only that unit was controlling the cursor. Now, let 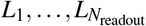 be the loss of the most dominant unit (*L*_1_) to the least dominant unit 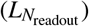 for a given seed and for a given value of CLDA intensity. The single-unit normalized reach performance was defined by

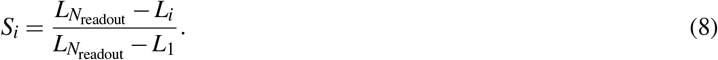

We thus have *S*_1_ = 1 and 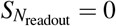.

A similar transformation was performed to obtain the normalized performance in Fig. 5F. If *L*_*i*_ is the loss after the *i*th most dominant unit has been added to the pool of readout units, then the normalized performance was

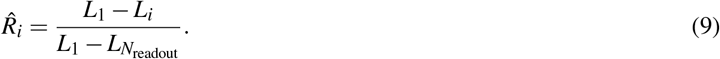

Here, we have 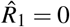 and 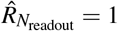.

Combinatorial unit-adding curves (Fig. 5G) were computed by sampling all combinations of size *s*, for *s* = 1, …, *N*_readout_, and evaluating the loss when each such combination was moving the cursor. To allow comparison across CLDA intensities, we normalized each loss using the minimum and maximum losses (across all subset sizes) when a fixed decoder was used:

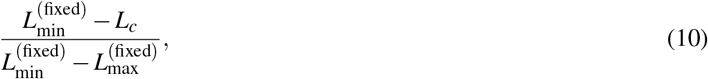

where *L*_*c*_ is the loss incurred for a specific combination.

The coefficients of determination in Fig. 5H were computed using the function `linregress` from the Python package `scipy.stats`. Each point in the scatter plots represents the change in a recurrent weight during learning with an adaptive decoder versus the weight change of the same connection with a fixed decoder, for a given network realization (random seed). Each plot contains 100^2^ weights for each of the 10 realizations.

