## supplementary materials for "Assistive sensory-motor perturbations influence learned neural representations"

#### Supplementary Methods

##### Neural tuning curves

Neural tuning curves were defined as the mean firing rate of the neuron in a time window (0-300ms after the go cue) as a function of the eight target directions. We first identified the mean firing rate of each unit per target direction and then modeled the relationship between neural activity and movement direction via a cosine tuning model [1]:

$$f = B_1 \cos\theta + B_2 \sin\theta + B_3$$

where $\theta$ represents the target direction and $B_1$, $B_2$ and $B_3$ are model coefficients. The coefficients were estimated via linear regression and then used to compute each unit's modulation depth (MD) and preferred direction (PD):

$$MD = \sqrt{B_1^2 + B_2^2}$$
$$PD = \text{arctan2}(B_2/B_1)$$

We calculated the change in tuning properties within each learning series by comparing MDs on early and late training days: $\Delta\text{MD} = \text{MD}_{\text{late}} - \text{MD}_{\text{early}}$. To compare these tuning changes to our model, we calculated the change in model coefficients between early and late training day: $\Delta\mathbf{w} = \mathbf{w}_{\text{late}} - \mathbf{w}_{\text{early}}$, where $\mathbf{w}_{\text{late,early}} \in \mathbb{R}^{n_{\text{units}}}$ were calculated from the model coefficients by taking the mean across time bins and then, for each unit, selecting the mean weight for the most contributing target direction. Figure S2B shows the correlation between $\Delta$MD and $\Delta\mathbf{w}$.

##### Comparison of offline velocity estimates

We used the same velocity Kalman filter decoders from the experiments to estimate cursor velocities offline. We compared velocity estimates obtained using all readouts and using only designated subsets of readouts (see below and Fig. S3G-I for detail). To quantify the similarity between the estimated velocities, we calculated the following time-average matching:

$$M = 1 - \frac{1}{T}\sum_{t=1}^{T} \begin{cases} 0 & \text{if } \|\mathbf{V}_t^{(\text{all})}\| = \|\mathbf{V}_t^{(\text{subset})}\| = 0 \\ \frac{\|\mathbf{V}_t^{(\text{all})} - \mathbf{V}_t^{(\text{subset})}\|^2}{\|\mathbf{V}_t^{(\text{all})}\|^2 + \|\mathbf{V}_t^{(\text{subset})}\|^2} & \text{otherwise} \end{cases}$$

where $\mathbf{V}_t^{(\text{all})}$ and $\mathbf{V}_t^{(\text{subset})}$ represent the cursor velocities estimated using all readouts and the subset, respectively. We defined the summand as zero when both velocities are zero. This ensures that $0 \leq M \leq 1$. We compared the velocities estimated using all readouts and the top $N_c^{\text{late}}$ units—from the target-encoding analysis—on late day (Fig. S3G). To assess how the addition of units improved the accuracy of cursor velocity estimates, we computed $M$ as we progressively added more units ranked according to the neuron adding curves obtained from the target-encoding analysis on early and late days (Fig. S3H). Finally, we compared the matching score $M$ for the two training phases when using a number of units equal to $N_c^{\text{late}}$ on both the early and late day (Fig. S3I).

#### Dimensionality estimation using Participation ratio

We computed the neural dimensionality for readouts and nonreadouts using the participation ratio (PR) [2, 3, 4]. PR summarizes the number of collective modes, or degrees of freedom, that the population's activity explores. It provides information about the spread of data and can be directly computed from the pairwise covariances among units $C_{\text{units}}$ in a population. It is defined by

$$\text{PR} = \frac{[\text{tr}(C_{\text{units}})]^2}{\text{tr}(C_{\text{units}}^2)}.$$

We estimated population dimensionality for readouts and nonreadouts separately each day. $\text{PR} \in [1, n_{\text{units}}]$, but since we had different number of nonreadout units that were recorded each day and different number of readout units across series, we normalized PR using

$$\text{PR}_{\text{norm}} = \frac{\text{PR} - 1}{n_{\text{units}} - 1}.$$

Therefore, $\text{PR}_{\text{norm}} \in [0, 1]$ with $\text{PR}_{\text{norm}} = 0$ when PR = 1 and $\text{PR}_{\text{norm}} = 1$ when $\text{PR} = n_{\text{units}}$. Note that a small difference in $\text{PR}_{\text{norm}}$ yields a big change in actual dimension (PR of 9.5 normalizes to $\text{PR}_{\text{norm}} = 0.25$ whereas PR of 11.2 normalizes to $\text{PR}_{\text{norm}} = 0.3$ for $n_{\text{units}} = 16$).

We estimated the dimensionality each day (Fig. S5A) using trials with completed reaches by creating a single time-series of concatenated spike counts from all reach segments (from go cue to the cursor entering the peripheral target, $n_{\text{units}} \times (n_{\text{timebins}} \times n_{\text{trials}})$. The readout population dimensionality was estimated using only the readout units used in the decoder that day. The change in dimensionality between early and late learning (Fig. S5C) was estimated by $\Delta\text{PR} = \text{PR}_{\text{late}} - \text{PR}_{\text{early}}$. We computed the 95% confidence intervals for PR calculations (Fig. S5C) by re-sampling the original trials with replacement, matching target identity distributions (N = $10^4$).

Dimensionality analysis for BCI learning with a fixed decoder (Fig. S5B, C) was performed on the dataset from [5]. It used the same method as above except that training epochs were used instead of days [6, 7]. Each consecutive training epoch contains a constant number of trials, which may combine data across training days. Note that our analyses differ from past studies [6, 7], which computed a measure of dimensionality based on the shared covariance extracted using factor analysis, instead of the total covariance as used by the PR metric.

#### Supplementary Tables

| Subject | # of readouts (early) | # of readouts (late) | # of readout units replaced | # of weight change events | # of ensemble change events | # of nonreadouts (early - late) |
|---|---|---|---|---|---|---|
| J | 16 | 16 | 1 | 4 | 1 | 36 - 36 |
| J | 16 | 16 | 4 | 3 | 1 | 39 - 39 |
| J | 16 | 16 | 3 | 4 | 2 | 41 - 41 |
| J | 16 | 16 | 2 | 2 | 1 | 38 - 38 |
| J | 16 | 15 | -2 +1 | 2 | 1 | 37 - 38 |
| J | 16 | 15 | -1 | 1 | 1 | 28 - 29 |
| J | 16 | 16 | 3 | 2 | 1 | 29 - 29 |
| S | 12 | 12 | 0 | 2 | - | 118 - 118 |
| S | 11 | 11 | 4 | 2 | 2 | 66 - 66 |
| S | 12 | 12 | 0 | 1 | - | 121 - 121 |

**Table 1.** Information about the number of readouts and nonreadouts, the number of readout units replaced during ensemble change events, the number of weight change events and the number of ensemble change events for every series analyzed in the main text. The '+' and '-' indicate the number of units newly added or removed, respectively. The '-' symbol in the ensemble change events column indicates no ensemble changes for that series. The range in the nonreadouts column denotes the total number of nonreadouts in the early and late stages.

### Supplementary Discussion

#### Compact Representation in Units Does Not Imply Compact Representation in Neural Modes

We observed compaction in both the “mode" space and the “individual unit" space. Here, we demonstrate that these observations do not simply follow from one another. We consider a two-class classification problem, eliminating the complexity of multiple targets, and we ignore the time dimension. We define $x$ as a random column vector representing centered single-unit activities and consider a logistic regression model $y(x) = f(w^\top x + b)$, where $^\top$ denotes transpose, and $w$ and $b$ are the model parameters. After fitting this model to data and achieving good generalization, we assume $w$ has been determined.

Drawing inspiration from Fig. 3, we define a representation as compact if the weight vector $w$ is sparse, specifically having a few dominant elements (ideally one, for the sake of this discussion). We introduce matrix $A$ containing the principal vectors as columns, forming an orthonormal matrix ($AA^\top = A^\top A = I$, where $I$ is the identity matrix). This allows us to reformulate the logistic regression model in terms of principal components and their projections: $y(x) = f((A^\top w)^\top (A^\top x) + b)$, where $A^\top x$ represents the principal components of $x$, and $A^\top w$ is the projection of $w$ onto the principal vectors.

If $w$ is sparse, its projection $A^\top w$ will effectively highlight the contribution of the dominant unit(s) in the space of principal vectors. However, whether the resulting weight vector in this transformed space remains compact (sparse) depends on the characteristics of the principal vectors and, thus, on the covariance matrix of the data. This observation underscores that a compact representation in “unit space" does not automatically imply a compact representation in “mode space”.

The essence of this discussion is that the transformation of the basis (through rotation or otherwise) does not necessarily preserve the sparsity of the representation. Thus, the compaction of representations across different spaces (unit vs. mode) is not a trivial matter and requires careful consideration of the underlying data structure and transformation methods used.

#### Impact of Neural Recording Methods

Differences in neural recording methods, such as threshold crossings versus sorted units, may also influence co-adaptation processes and the stability of neural representations. For instance, we used threshold crossings in monkey J. Threshold crossings may have different stability properties over days than sorted units, which might lead to differences across monkeys. However, isolating contributions of neural recording properties is beyond the scope of what is feasible with these current data. Future studies exploring how neural recording stability, specifically, influences neural plasticity and decoder adaptation algorithms will be critical for designing BCIs that maintain performance over time.

#### Supplementary Figures

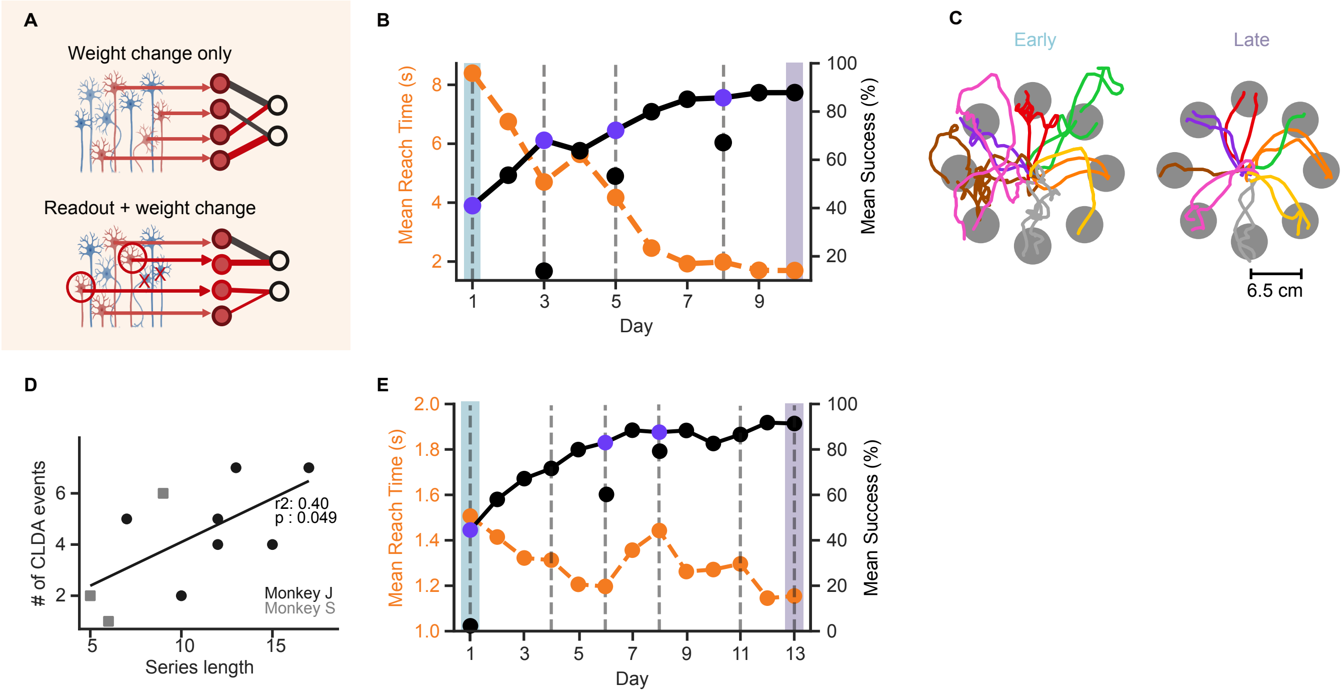


**Figure S1. Supplementary figure related to figure 1: example behavior from monkey S and J (A)** BCI decoder adaptation. During learning, the decoder was adapted in two ways: changing the weights relating readout units to velocity while keeping the identify of readout units constant (Weight change only), and altering both the contributing readout units and their corresponding decoder weights (Readout + weight change). **(B)** Success percentage (solid black line) and mean reach time (dashed orange line) across days for an example learning series for monkey S (seba010911_011811). Indigo dots show performance boost after decoder adaptation. Early (blue) and late (purple) training phases are indicated. Vertical dashed lines indicate days where decoder was adapted - gray dashed lines (weight change only) and black dashed lines (readout + weight change). **(C)** Cursor trajectories during early (left) and late (right) training phases for the example series in panel B. **(D)** Number of total decoder adaptation events (including both weight changes and readouts + weight changes) plotted against series length for monkey J (black circles) and monkey S (grey squares). **(E)** Same as (B) for another example series from monkey J where decoder was adapted on day 1 for better performance.

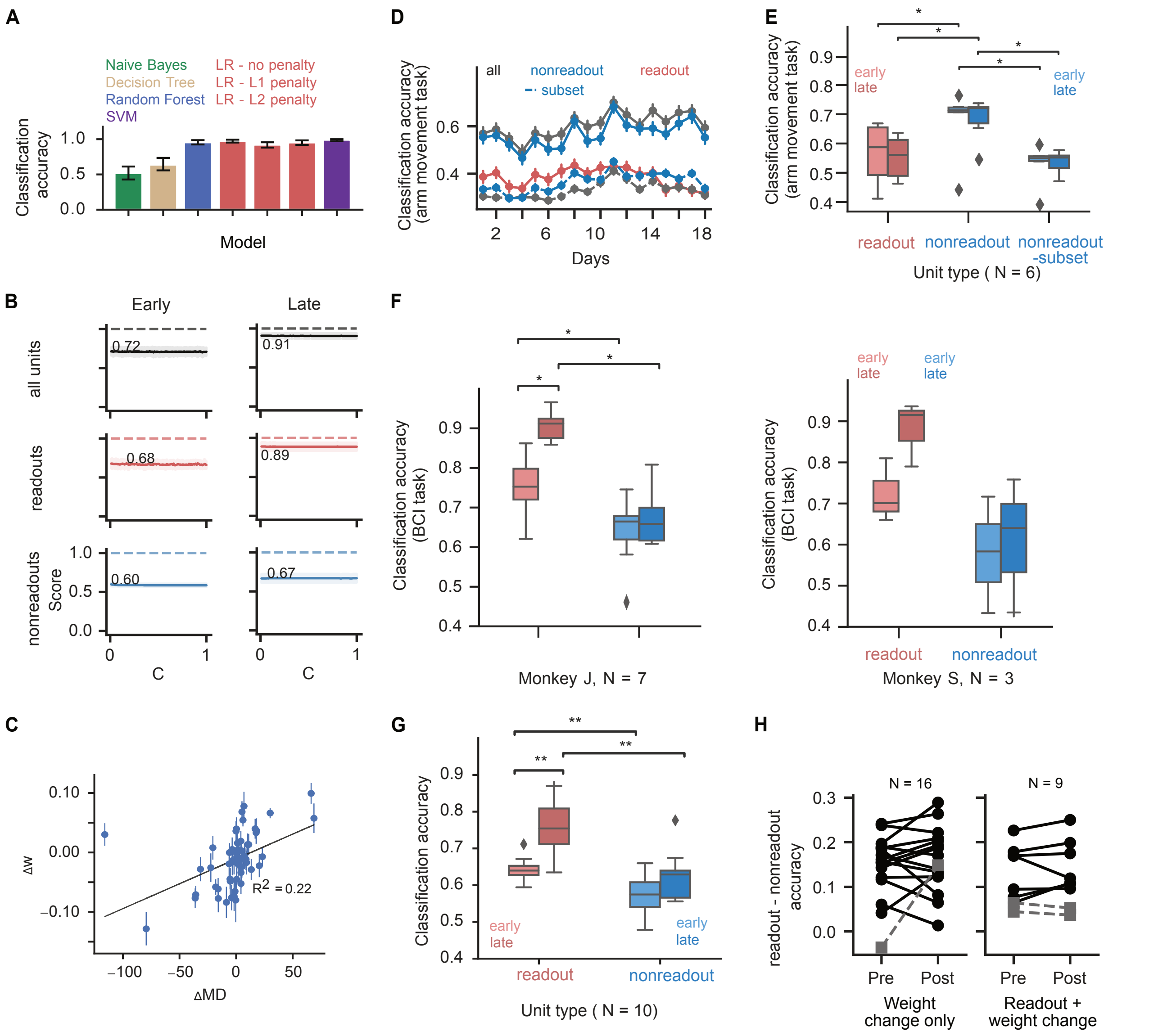


**Figure S2. Supplementary figure related to figure 2: credit assignment** (A) Classification accuracy obtained from different classifier models when trained on neural data from last day of our example series. Error bars are 95% confidence intervals on test accuracy. (B) Early and late classification scores for all units (black), readouts (red) and nonreadouts(blue) plotted as hyperparameter $C$ (related to regularization strength) varies from $10^{-5}$ to 1. (C) Change in logistic regression model coefficients ($\Delta \mathbf{w}$) between early and late day as a function of changes in modulation depth of units from example series. (D) Classification analysis from all units (black), readouts (red) and nonreadouts (blue) for arm movement task for the example series. There were significantly more units in the 'all' and 'nonreadout' populations compared to the 'readout' population, which contributes to the large difference in overall classification accuracy. Dashed blue and black lines show the classification accuracy achieved from a subset of nonreadout and all units, respectively, randomly drawn to match the number of readout units. (E) Early vs late classification accuracy for readout (shades of red) and non-readout populations(shades of blue) across all series for arm movement task (readout-early vs nonreadout-early: $^*p = 0.031$; readout-late vs nonreadout-late: $^*p = 0.031$; nonreadout-early vs nonreadout subset-early: $^*p = 0.031$ ; nonreadout-late vs nonreadout-late subset: $^*p = 0.031$, readout-early vs nonreadout-subset early: $p = 0.2$ (n.s); readout-late vs nonreadout-subset late: $p = 0.8$ (n.s), $n = 6$, Wilcoxon signed-rank test). (F) Early vs late classification accuracy for readout (shades of red) and nonreadout populations (shades of blue) shown separately for each monkey for BCI task. Monkey J ($n = 7$) : nonreadout-early vs. nonreadout-late: $p = 0.3$ (n.s); readout-early vs. readout-late: $^*p = 0.016$; readout-late vs. nonreadout-late: $^*p = 0.016$; readout-early vs. nonreadout-early: $^*p = 0.016$. Monkey S ($n = 3$): nonreadout-early vs. nonreadout-late: $p = 0.5$ (n.s); readout-early vs. readout-late: $p = 0.25$ (n.s); readout-late vs. nonreadout-late: $p = 0.25$ (n.s) ; readout-early vs. nonreadout-early: $p = 0.25$ (n.s). Wilcoxon signed-rank test. (G) Early vs late classification accuracy for readout (shades of red) and nonreadout populations (shades of blue) from BCI task using neural activity from the go cue to 200 ms after go cue. Nonreadout-early vs. nonreadout-late: $p = 0.1$ (n.s); readout-early vs. readout-late: $^{**}p = 0.008$; readout-late vs. nonreadout-late: $^{**}p = 0.004$; readout-early vs. nonreadout-early: $^{**}p = 0.008$. Two-sided Wilcoxon signed-rank test, $n = 10$. (H) Readout-nonreadout classification accuracy, grouped by the type of change (with (right) and without (left) readout unit changes) across all series and monkeys. Weight change only: $p = 0.3$ (ns), $n = 16$; readout + weight change: $p = 0.25$ (ns), $n = 9$. Pre < post, one-sided Wilcoxon signed-rank test.

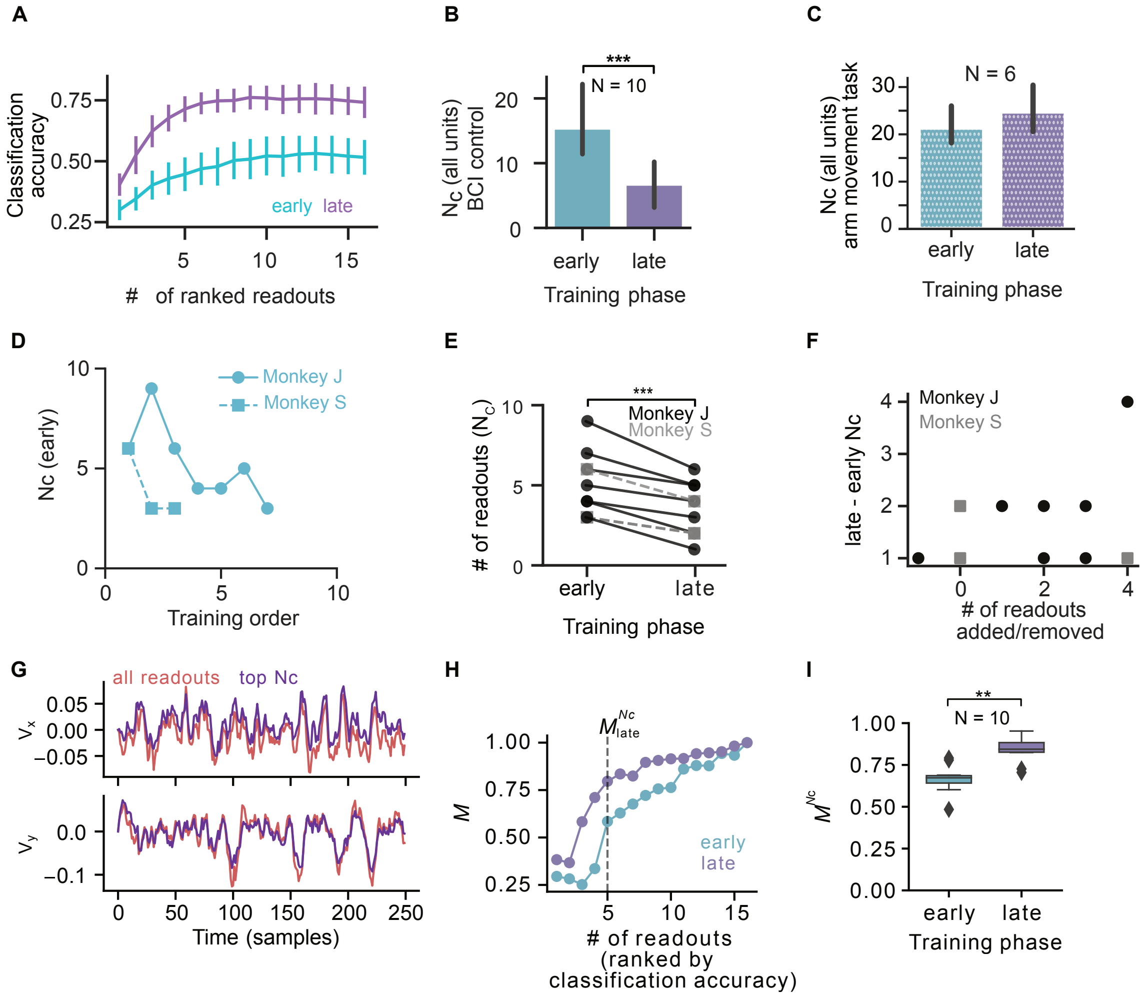


**Figure S3. Supplementary figure related to figure 3: compaction** (A) Classification accuracy (not normalized) as units are added in ranked order (ranked NAC) on early (cyan) vs late (purple) day for the representative series (Unnormalized version of Fig 3F). (B) Comparison of $N_c^{\text{late}}$ vs $N_c^{\text{early}}$ across series while using all units ($N_c^{\text{late}}$ is less than $N_c^{\text{early}}$, $^{***}p = 9.8 \times 10^{-4}$, $n = 10$, Wilcoxon signed-rank test ). (C) Similar to (B) for arm movement task ($p = 0.07(n.s)$ , $n = 6$, Wilcoxon signed-rank test). (D) $N_c^{\text{early}}$ for readouts across chronologically-ordered series for monkey J (solid lines, round markers) and monkey S (dashed lines, square markers). (E) Comparison of $N_c^{\text{late}}$ and $N_c^{\text{early}}$ across series ($N_c^{\text{late}} < N_c^{\text{early}}$, $^{***}p = 9.8 \times 10^{-4}$, $n = 10$, one sided Wilcoxon signed-rank test). Monkey J (black circles); Monkey S (grey squares).. (F) Late−early $N_c$ vs. number of readouts added or removed in a learning series ($R^2 = 0.36$, $p = 0.3$ (n.s). (G) X and Y cursor velocities reconstructed offline using the Kalman filter decoder with all readouts (red) and the top $N_c^{\text{late}}$ readout units (purple) identified by target classification analysis for the representative series. (H) Matching $(M)$ between the velocities reconstructed using all units vs. as units are added in ranked order (ranked NAC) on early (cyan) vs late (purple) day for the representative series. (I) Velocity matching ($M^{N_c}$) when using top Nc units on early vs late day ($^{**}p = 0.0019$, $n = 10$, Wilcoxon signed-rank test).

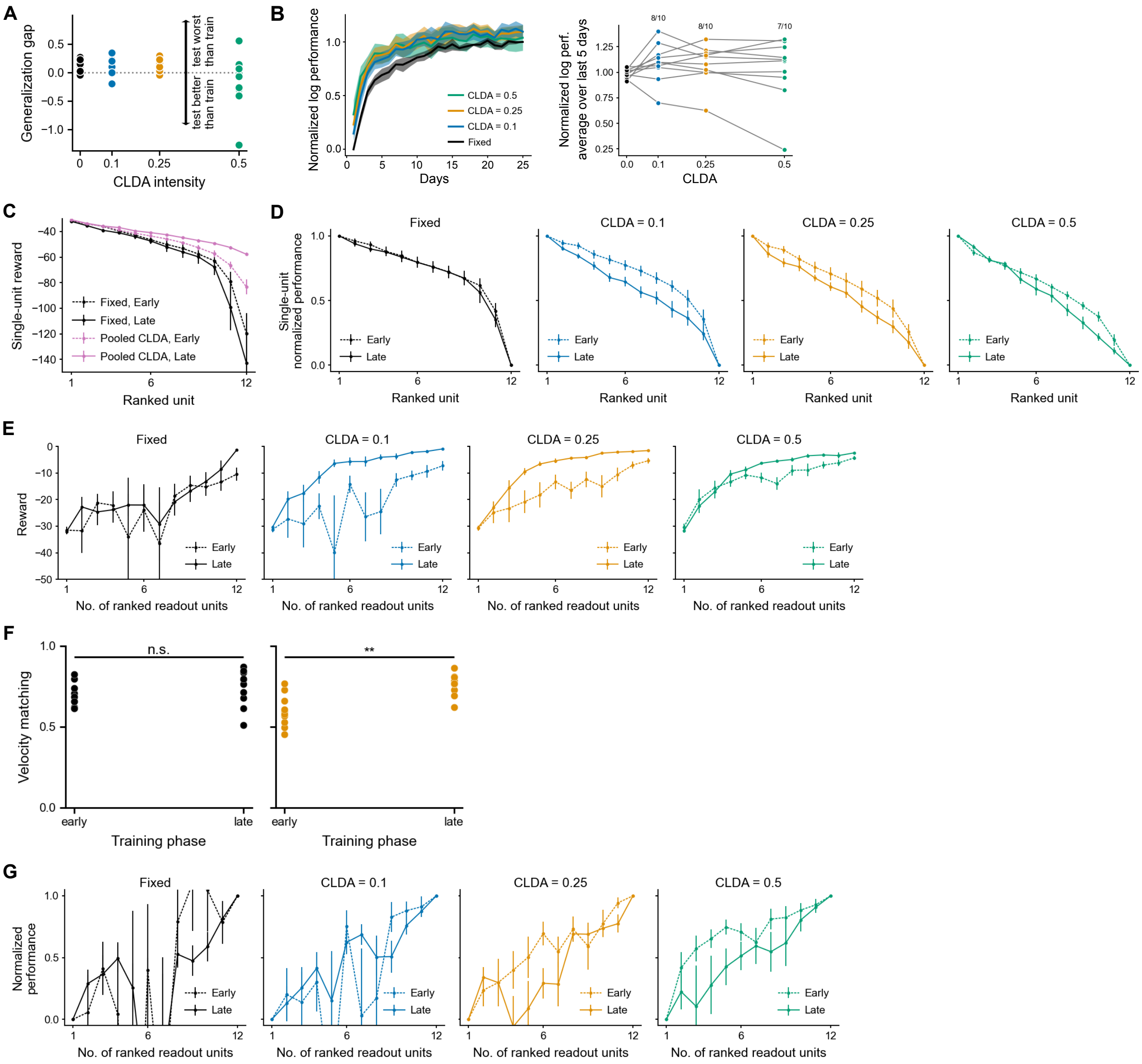


**Figure S4. Supplementary figure related to Fig. 4: model.** (A) Generalization gap is performance on training set minus performance on test set. Gap for the adaptive decoder is not significantly different from that for the fixed decoder ($p > 0.1$, $n = 10$, Wilcoxon signed-rank test, CLDA $>$ fixed). Variance for CLDA is also not larger than variance for the fixed decoder ($p > 0.1$, $n = 10$, paired permutation test for variance(CLDA$>$ 0) $>$ variance(fixed)) (B) Training performance when stopping CLDA after a loss criterion was reached (i.e., when loss $< 2$). (C-D) Unnormalized pooled (C) and normalized un-pooled (D) single-unit performances using the reward which is the negative of the loss. (E) Unnormalized NACs. (F) Matching between the online velocity inferred using the top 6 best units—according to the loss-based performance metric (see Fig. 4F)—and using all readout units. Left: fixed decoder. Right: adaptive decoder with CLDA = 0.25, which produced the most salient compaction according to Fig. 4F ($p = 0.002$ (**), $n = 10$, Wilcoxon signed-rank test). (G) Compaction results when stopping CLDA.

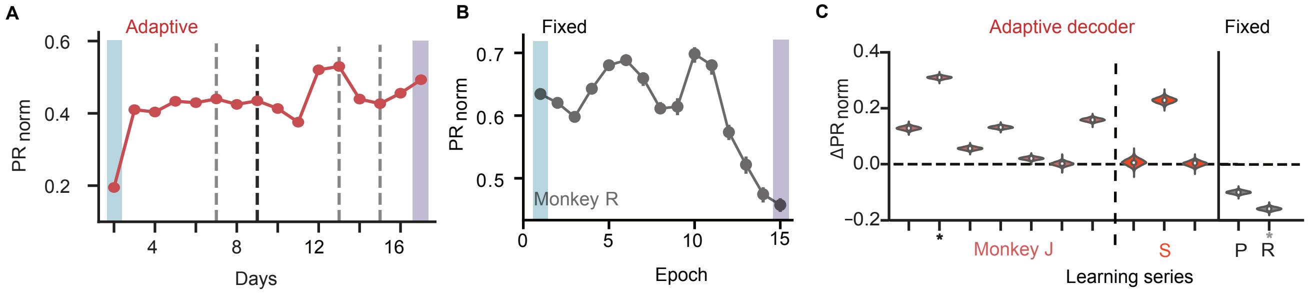


**Figure S5. Supplementary figure related to Fig. 5: Population analysis** (A) Normalized participation ratio ($PR_{norm}$) for an example adaptive decoder learning series (same as in Fig. 1D). Error bars are 95% confidence intervals estimated via bootstrapping (see Methods). Note that most error bars are hidden by data point markers. (B) Same as (A) for fixed decoders (monkey R; data from [5]). Note: the x-axis denotes epoch, which are groups of constant number of trials, consistent with previous analyses [6]. (C) $\Delta PR_{norm}$ distributions for learning series with adaptive decoders ($n = 7$ for monkey J and $n = 3$ for monkey S). Fixed data is from two monkeys (P and R) using fixed decoders (data from [5]). * denotes the example series shown in panels A and B.
